# Oligonucleotide genetics for *Pseudomonas aeruginosa* enables high throughput hypomorph screening

**DOI:** 10.64898/2026.05.19.726351

**Authors:** Sundar Pandey, Ayesha M. Ahmed, Kanna Nagamatsu, Melissa Reyes, Jiwoong Kim, Xiaowei Zhan, David E. Greenberg, Scott H. Saunders

## Abstract

The continual advancement of genetic tools has been critical to our modern understanding of bacteria, with transposons, plasmids, and homologous recombination becoming workhorses of molecular microbiology. However, precisely specified reverse genetic approaches remain painstakingly slow and inaccessible, particularly in non-model strains. This reality is exemplified by the opportunistic pathogen, *Pseudomonas aeruginosa* (Pa), where conventional allelic exchange remains the dominant reverse genetic method. Here, we adapt a rapid genetic toolkit for use in Pa, relying directly on commercially available oligonucleotides (120 bases) to create precise genomic mutations through homologous recombination (i.e. oligo recombineering). Oligo Recombineering followed by Bxb-1 Integrase Targeting (ORBIT) uses a short attachment site for an integrating plasmid, which provides traditional antibiotic selection and can also carry flexible cargo. We establish Pa ORBIT works effectively for gene deletion without off target mutations, optimize protocol parameters (e.g. oligo length, electroporation), and demonstrate markerless and clean deletions. Importantly, our toolkit works well in clinical Pa strains as demonstrated by constructing efflux pump deletions in three different isolates. To test the high throughput capabilities of Pa ORBIT, we created over 160 degron-based hypomorphs (i.e. knockdowns) across 43 essential proteins in a pooled mutant library. Upon screening this library with and without antibiotics, we identify highly vulnerable essential proteins and hypomorphs that display synergy with clinical drugs. Therefore, ORBIT can be used for cutting edge low and high throughput investigations in this priority pathogen, setting the stage for answering critical basic and clinical science questions.

**Significance:** To understand bacterial genomes, researchers need access to rapid, flexible, precise and high throughput genetic perturbation tools. Here we present an oligonucleotide-based method that satisfies these requirements for use in the opportunistic pathogen, *Pseudomonas aeruginosa*. By relying on oligos to encode genomic homology arms, no molecular cloning is required – making these tools rapid, robust, and scalable. We benchmark gene deletions in both lab and clinical strains, opening the possibility of rapid genetic studies across the *P. aeruginosa* pangenomic space. At high throughput, we use an oligo pool to create a mutant library of degron tagged essential proteins. These knockdowns (i.e. hypomorphs) show certain essential genes are highly vulnerable and others are synergistic with clinical drugs, providing insight into future antibiotic and co-therapy development.

## Introduction

The opportunistic bacterium, *Pseudomonas aeruginosa* (Pa), causes a range of clinical infections, often in hospitalized or immunocompromised patients, and treatment of these infections is becoming increasingly difficult due to both endogenous and acquired antimicrobial resistance. To combat this important pathogen, fundamental research is needed to better understand its complex basic biology and clinically relevant phenotypes. There is a rich history of using molecular genetic approaches to directly perturb genotypes and assess causation in bacteria and Pa specifically. For decades, transposons have played an important role in “forward” bacterial genetics, enabling high throughput selections, arrayed mutant libraries (1), and now complex pooled assays via Tn-Seq (2, 3). For targeted “reverse” genetics in Pa, allelic exchange using a time-consuming multi-step cloning procedure remains the dominant method (4, 5). Despite some success with newer genetic technologies (e.g. CRISPRi) (6–8), it remains unclear how to implement reverse genetics that is flexible, fast and high throughput in Pa.

Modifying the genome by directly introducing chemically synthesized DNA oligonucleotides into cells, oligo recombineering, has become a powerful approach to understand and engineer bacteria, particularly model *E. coli* (9–11). Oligo incorporation is catalyzed by single stranded DNA annealing proteins (SSAPs) that facilitate binding of homologous oligo and genomic sequences. SSAPs are known to be relatively host specific through interactions with endogenous single stranded DNA binding protein (SSB), and functional SSAPs have recently been found for diverse bacterial species, including Pa (12, 13). This oligo-based approach offers advantages in speed and throughput, since it requires no molecular cloning, however, it has typically been limited to very small mutations (1-5 base changes) that can be recovered without selectable markers, which limits its utility.

To overcome the limitations of oligo recombineering, Murphy et al. established a strategy that uses a short attachment (att) site on an oligo to direct the integration of a plasmid, providing a way to couple an oligo specified modification to plasmid encoded selectable markers (14). We recently adapted this method, Oligo Recombineering followed by Bxb-1 Integrase Targeting (ORBIT), for use in *E. coli* (15), and showed that it is highly effective for constructing varied mutations at low and high throughput. Importantly, this strategy enables the use of commercially available oligo pools to flexibly specify libraries of mutations. Because oligo pools are now available at extremely large scale, we were able to construct precisely specified mutant libraries containing over 100k members in *E. coli* to ask sophisticated genome wide questions (16). Here, we adapt ORBIT for use in Pa, optimize parameters and share protocols and plasmids to facilitate adoption by the Pa research community. We demonstrate ORBIT’s utility in drug resistant clinical Pa strains to knock out antibiotic efflux pumps and implement a genome wide mutant library.

An important genome wide question for Pa is to understand which essential gene products are easiest to inhibit and therefore promising drug targets to pursue. Using CRISPRi knockdowns in *Mycobacterium tuberculosis*, Bosch et al. ranked essential proteins by “vulnerability” (i.e. sensitivity to knockdown), and they found that some leading clinical drugs do not target particularly vulnerable proteins and suggested re-prioritizing antibiotic development efforts (17). Furthermore, the use of essential gene knockdowns, also called hypomorphs, has proved valuable in the discovery of novel antibiotics. For both established molecules and novel drug candidates, Johnson et al. found that hypomorphs of target (or related) proteins show differences in sensitivity in pooled mutant libraries, offering critical mechanistic information and a broader range of active molecular scaffolds for drug development (18, 19). However, constructing protein level knockdown hypomorphs has been a painstaking process, limiting wider adoption. In this work, we develop a degron based toolkit for essential gene knockdown and apply it at high throughput to characterize gene vulnerability and drug synergy.

One novel antibacterial strategy is sequence-based inhibition using short antisense oligos that penetrate bacterial cells and inhibit specific essential genes. Nonstandard oligonucleotides (11 bases) are coupled to cell-penetrating peptides, base-pair with target mRNAs, and block translation (20, 21). We have successfully used these peptide-conjugated phosphorodiamidate morpholino oligomers (PPMOs) as preclinical inhibitors of a variety of bacterial pathogens, targeting different essential genes (22–25). However, it remains inefficient to synthesize and screen PPMOs for activity, and new genetic screening techniques are needed to prioritize genes for targeted PPMO optimization. Here, we compare high throughput knockdown vulnerability to PPMO inhibition and find a synergistic PPMO-levofloxacin effect.

In this work, we demonstrate our effective ORBIT system for Pa at both low and high throughput and investigate the vulnerability of essential protein targets. We find notable variation in the vulnerability of Pa proteins and identify a range of interesting targets (e.g. Fis, NrdA). Given the flexibility and effectiveness of Pa ORBIT, we believe it will be a valuable method for investigating many important genetic questions in Pa and other important bacteria.

## Results

### Establishing ORBIT tools for *P. aeruginosa*

The goal of ORBIT is to create a genetic modification specified with a DNA oligo. This is accomplished by including a short attachment site on the oligo (attB) that then serves as an integration site for a non-replicating plasmid. Upon integration, the plasmid selectable marker enables the recovery of recombinant cells that possess the oligo-specified mutation. Therefore, ORBIT is comprised of two sequential molecular steps 1) RecT mediated oligo recombineering and 2) Bxb-1 mediated plasmid integration. To create an effective ORBIT system for *P. aeruginosa*, we first had to construct a replicating helper plasmid encoding RecT and Bxb-1. Single stranded annealing proteins used for oligo recombineering are known to be host specific, so we used PapRecT and an m-toluic acid inducible expression system, which was shown to effectively incorporate SNPs via oligo recombineering in Pa (Fig 1A) (12). Bxb-1 is known to be highly active across hosts, so we used the same arabinose inducible system as ORBIT for *E. coli*. Because *E. coli* integrating plasmids already do not replicate in Pa, we decided to continue using R6k Ori pInts, which will let ORBIT tools be shared easily across species. Throughout this work we refer to the two required ORBIT plasmids as “pHelper” (replicating with helper functions) and “pInt” (non-replicating, integrates with selectable marker).

**Figure 1.**
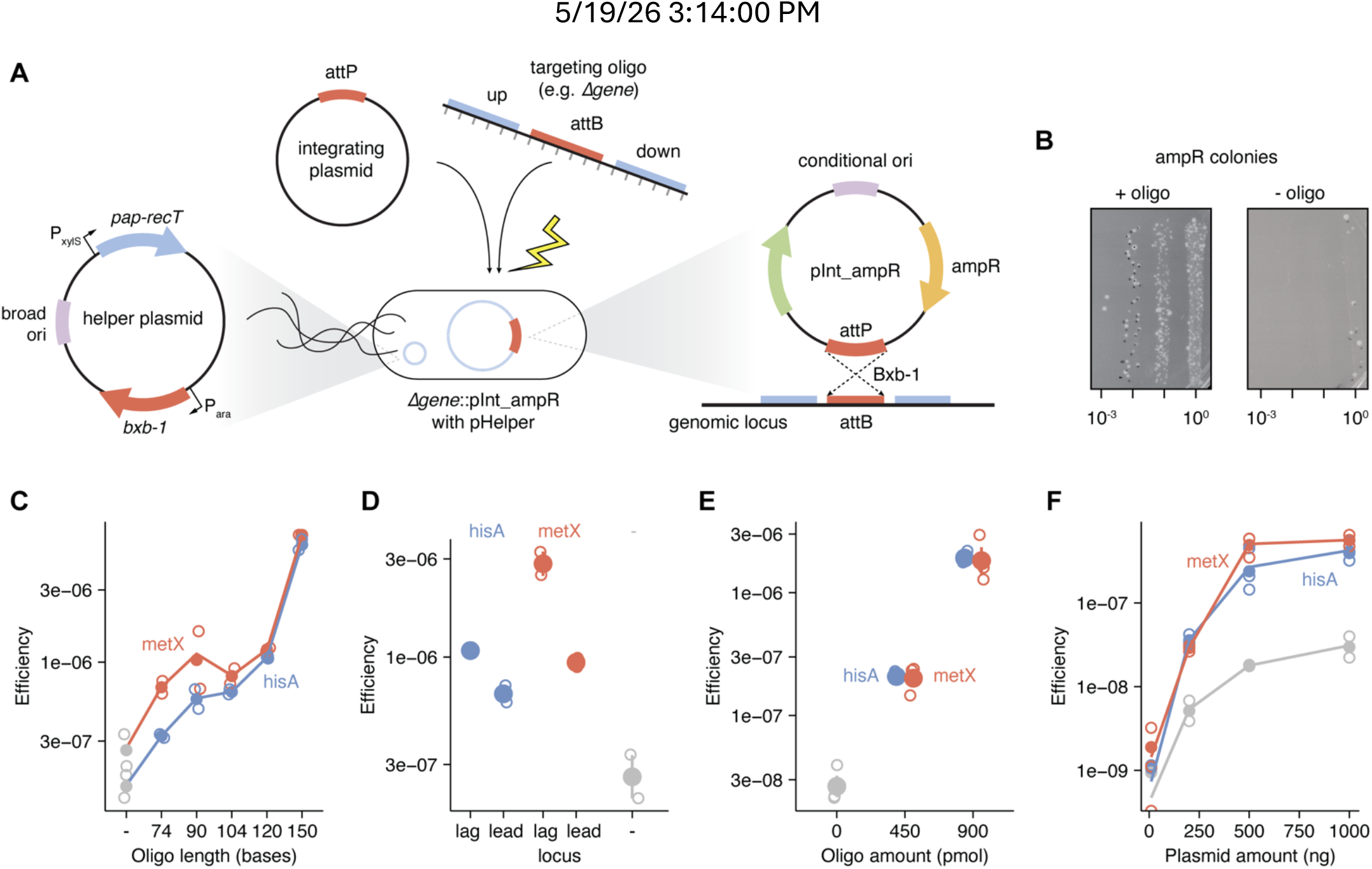
Pa ORBIT principle and optimization. A) Diagram of ORBIT helper plasmid along with oligo and integrating plasmid co-electroporation. B) ORBIT with Δ*hisA* targeting oligo yields antibiotic resistant colony forming units. C-F) ORBIT efficiency (i.e. ampR colonies / all colonies) for protocol variables (n=2 transformations, open circles. Solid points show mean values and bars show standard error). Dash denotes a negative control without oligo. Variables include oligo length (C), oligos targeting lagging strand and leading strand template (D), oligo amount (E), and integrating plasmid amount (F).

After combining the separately inducible ORBIT functions into a broad host range helper plasmid, we performed several rounds of plasmid troubleshooting and optimization before achieving consistent ORBIT modifications in model PAO1 and PA14 strains (26, 27). Following our initial development stage with maximal quantities of oligo and plasmid, we routinely observed that co-electroporation with a 120 nucleotide (nt) oligo (41 nt homology arms + 38 nt attB) and carbenicillin resistant integrating plasmid (i.e. pInt_ampR) yielded significantly more antibiotic-resistant colonies than transforming the plasmid alone (Fig. 1B), and these initial mutants were confirmed to have the correct locus structure by PCR.

Next, we used two gene deletions, Δ*hisA* and Δ*metX* (histidine and methionine biosynthesis respectively), to optimize the efficiency (i.e. recombinant frequency) of the protocol parameters. In PA14, we found that longer oligos were more efficient, with 150 nt oligos yielding 5.6-fold higher efficiency than 120 nt oligos (Fig. 1C). However, due to the dramatic increase in price, we typically used 120 nt oligos with consistent success. Oligo recombineering generally followed the expected trend established in other organisms, where oligos targeting the lagging strand template were more efficient than those targeting the leading strand template (Fig. 1D). However, this difference was not as striking as in *E. coli*, suggesting there may be differences in the oligo recombineering mechanism (11). Next, we titrated the amount of oligo and found that doubling the oligo amount to 900 pmol yielded ten times higher efficiency (Fig. 1E). Then we titrated the amount of integrating plasmid and found that efficiency saturates above 500 ng, and the off-target rate increases proportional to the on-target rate (Fig.1F). We tested a common strategy to limit oligo degradation, phosphorothioate bonds, but we observed no effect (Fig. S1A). We found the optimal recovery time following electroporation was overnight, although shorter recoveries also worked (Fig. S1B). Finally, we tested the effect of polymyxin B nonapeptide (PMBN) treatment, which was reported to improve transformation efficiency (28). Our results showed a 10-fold increase in ORBIT efficiency at the highest dose (Fig. S1C), but we did not regularly use PMBN due to a slower growth rate and higher cost. This work led us to a standard co-electroporation protocol for routine PA ORBIT that uses unmodified 120 nt oligos, 300 ng of freshly prepared integrating plasmid, and an overnight recovery (see supplementary information).

### Benchmarking gene deletion efficiency and accuracy

To benchmark the functionality of the ORBIT system in *P. aeruginosa*, we rigorously assessed gene deletions of our test loci *hisA, metX*, as well as the protease *clpP*. Then we employed ORBIT more broadly to construct gene deletions at other loci of interest (Fig. 2A). Across triplicate experiments in strain PA14 with *hisA, metX*, and *clpP*, we observed efficiencies ranging from 1 × 10^-6^ to 4 × 10^-6^, with Δ*clpP* being the most efficient target (Fig. 2B). This corresponded to a yield of hundreds to thousands of putative recombinant colonies from each transformation. PCR verification confirmed precise integration events at each locus, yielding gel-based accuracy from 37 to 100% (Fig. 2C, S2). The expected amplicon sizes were consistent with successful gene replacement with pInt_ampR, and products were Sanger sequenced to confirm perfect matches to the expected locus sequence (Fig. 2D, S2, S3). Expected auxotrophic mutants, Δ*hisA* and Δ*metX* did not grow on minimal medium, unless supplemented with histidine or methionine, respectively (Fig. 2E, S2D). ClpP mutants are known to exhibit growth defects at high temperatures, and we observed strong decreases in CFUs at 42°C compared to wild type PA14 (Fig. S2E). Therefore, each ORBIT test mutation was confirmed at the molecular and phenotypic level.

**Figure 2.**
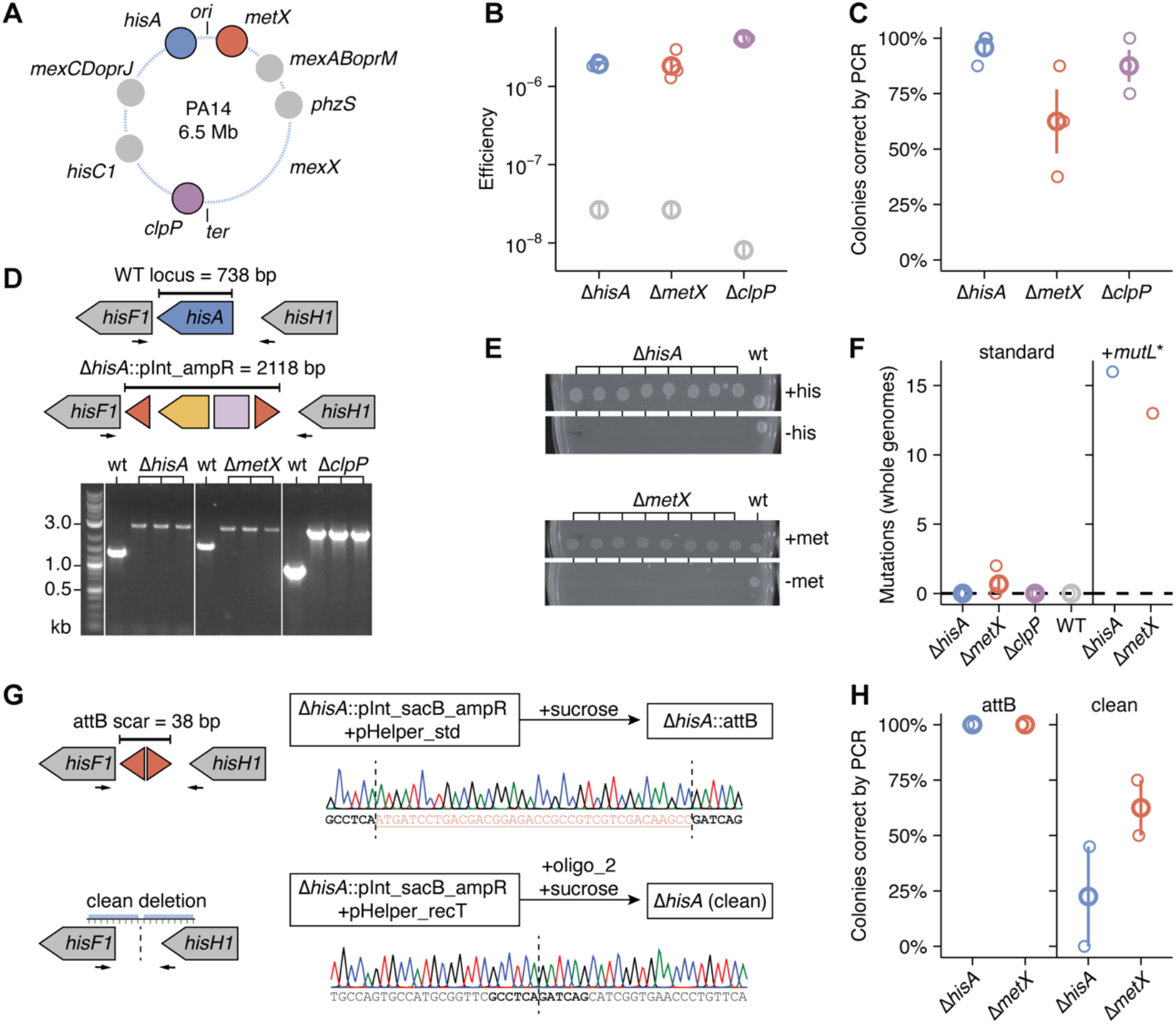
Gene deletion benchmarks and markerless modifications. A) Genes successfully deleted by ORBIT. B)Efficiency of gene deletions (n=3 transformations), including no oligo controls (gray dots). C) Accuracy of gene deletions by PCR. D) Representative PCR results spanning each ORBIT modification. E) Representative validation for ΔhisA::pInt_ampR and ΔmetX::pInt_ampR (n=8 colonies shown) on minimal medium with or without the expected auxotrophic amino acid. F) Whole genome sequencing results for ORBIT modified strains. Total number of off target mutations are reported relative to WT. For standard ORBIT ΔhisA, ΔmetX, and ΔclpP, n=3 independent colony whole genome sequences. For WT and +mutL strains, n=1 whole genome sequence. G) Markerless ORBIT modification achieved through pInt_sacB_ampR counterselection to generate 38 bp attB scar (top) or clean deletion using a second oligo without attB (bottom). Clean deletion is achieved with a helper plasmid only containing PapRecT, not Bxb-1. H) Accuracy of markerless modifications shown in G (by PCR, n=2 independent transformations).

Next, we sought to test our recombinant strains for off target single nucleotide polymorphisms (SNPs) accumulated in the genome. Importantly, our ORBIT helper plasmid does not include any mismatch repair inhibition mechanism, which is commonly used to increase efficiency of traditional oligo recombineering. For 8 of the 9 constructed strains, we detected zero off target SNPs relative to the parental strain, while a single Δ*metX* strain harbored two SNPs (Fig. 2F, S2F). In contrast, using an earlier ORBIT helper plasmid with the PaMutL* allele (mismatch repair inhibition), we observed >10 mutations throughout the genome for multiple strains. Generally, strain PAO1 yielded slightly lower efficiency, but very similar accuracy and SNP results across the *hisA, metX* and *clpP* loci.

Beyond our three test loci, we also performed 9 other deletions across the PA14 genome. ORBIT proved highly reliable and multiple correct colonies were rapidly obtained, verified by Sanger sequencing, and showed the expected phenotypes (e.g. Δ*phzS* loses pyocyanin pigmentation). However, we did observe variation in efficiency and accuracy at different loci, which we expect was caused by a combination of genomic factors, oligo properties, and daily competent cell variability. Together, these results establish the successful implementation of ORBIT in *P. aeruginosa*, enabling precise, reproducible chromosomal deletions with essentially no off-target mutations using commercially available DNA oligonucleotides. This method is significantly faster than standard allelic exchange, requiring no cloning and 1 day of hands on work (protocol spans 4 days including pre-culture, overnight recovery and plate incubation) to obtain initial recombinants (5). To facilitate the adoption of ORBIT by the Pa community, we will release plasmids on Addgene, provide protocols as supplemental material and established oligo design tools for PAO1 and PA14 model genomes (see GitHub repository).

### Marker-less modifications and large integrating plasmids

To generate marker-less and scarless deletions in Pa, we had to contend with a previously observed feature of Bxb-1 — in addition to the integration reaction, the opposite excision reaction also occurs at low rates (15). We exploited this reversible property to rapidly create marker-less mutants. Using our standard Pa helper plasmid, we performed ORBIT with a Δ*hisA* targeting oligo and an integrating plasmid containing the counter selection marker *sacB* (also on pHelper). Following successful integration, recombinants (still with pHelper) were simply plated on sucrose, which is toxic in the presence of *sacB*. By PCR and Sanger sequencing, 100% of tested colonies successfully excised the ORBIT integration, leaving only the 38 nt attB site as a scar (Fig 2G). Given that the attB scar would be active upon sequential ORBIT modification, we also confirmed that FLP based excision can be used to create larger, inert scars (Fig. S4A)(29). Next, we tested alternative att sites that contain different central dinucleotides (attBP1 ‘GT’; attBP2 ‘TT’; attBP5 ‘CT’), which show strong orthogonality in *E. coli* (15). The variant att site 2 showed high efficiency and accuracy (Fig. S4B-D), raising the possibility of creating attBP2 ORBIT modifications in markerless strains that possess attB1 scars.

However, the Pa community routinely prefers “clean” or scarless deletions given that sequential mutations can be confidently made in these backgrounds. To achieve scarless deletion we took the same approach as in *E. coli*, where an initial ORBIT modification uses an integrating plasmid with *sacB* and then a second oligo cleanly deletes the pInt_sacB integration. Unfortunately, likely due to rates of Bxb-1 excision exceeding ORBIT deletion efficiencies, we were rarely able to obtain scarless mutants in the presence of Bxb-1. Excision of pInt, leaving an attB scar, was the predominant genotype. This was true even after adopting a rhamnose inducible system with very low leaky expression of Bxb-1 (Fig. S5A-C)(30). Therefore, we constructed a helper plasmid that contained only PapRecT and not Bxb-1 for use during this step. We also needed to be able to remove the original pHelper without sucrose counter selection, so we introduced a temperature-sensitive origin of replication to pHelper (Fig. S5D-E)(31). This plasmid worked effectively for ORBIT and loss of the plasmid was confirmed by PCR following incubation at 42 °C, with nearly complete curing observed within a single passage. Beginning from Δ*hisA*::pInt_sacB_ampR and Δ*metX*::pInt_sacB_ampR strains carrying the pHelper with only PapRecT, we successfully constructed scarless deletion mutants upon electroporation with a clean deletion oligo and counter selection on sucrose (Fig. 2GH). Further work will be required to streamline the clean deletion protocol, as this process requires intermediate curing of the Bxb-1 containing temperature sensitive pHelper, and subsequent transformation with the pHelper with only PapRecT. However, this tool should be valuable at present since it requires no molecular cloning to achieve – just two oligos.

In contrast to deletion, we also frequently use ORBIT to stably complement or place reporters onto the genome. To understand the size limits of integrating plasmids, we attempted to integrate pInts carrying a range of cargo sizes into the *clpP* locus. We successfully integrated the entire violacein biosynthetic pathway under the control of the luxR regulator (8.7kb cargo, 11 kb total), as verified by PCR and observed purple pigmentation (Fig. S5F-I)(32). This demonstrates that ORBIT is a flexible tool for genome manipulation, enabling precise deletion and large construct insertion.

### Rapid genetics in drug-resistant *P. aeruginosa* clinical isolates

It remains difficult to perform genetic manipulation across bacterial strains due to high phenotypic (e.g. permeability & growth rate) and genotypic (e.g. defense systems) variability. Further, different genomic sequences typically require separately cloned constructs; however, ORBIT relies on short homology that is less likely to vary and oligos that are easily customized to each strain. To assess the utility of ORBIT in genetically diverse Pa backgrounds, we started with a set of ten clinical isolates from the Texas Infectious Diseases Biorepository (TIDB) that were known to be relatively gentamicin and ampicillin sensitive, as required for our current plasmids. To start, we tried transforming the helper plasmid and confirmed its replication by native plasmid miniprep. Only three strains grew effectively and maintained the helper plasmid: TIDB3146, 3301, and 4041—representing distinct infection sources, sequence types, and accessory genomes (Fig. 3A-B). The untransformed strains showed diverse phenotypic properties like slow growth rates and mucoidy, and they may also possess defense mechanisms like CRISPR or restriction-modification systems that inhibit helper plasmid transformation. The three pHelper-transformable isolates displayed variable resistance to β-lactams and fluoroquinolones, including ciprofloxacin resistance in isolates 3146 and 3301, and high piperacillin-tazobactam resistance in isolate 4041.

**Figure 3.**
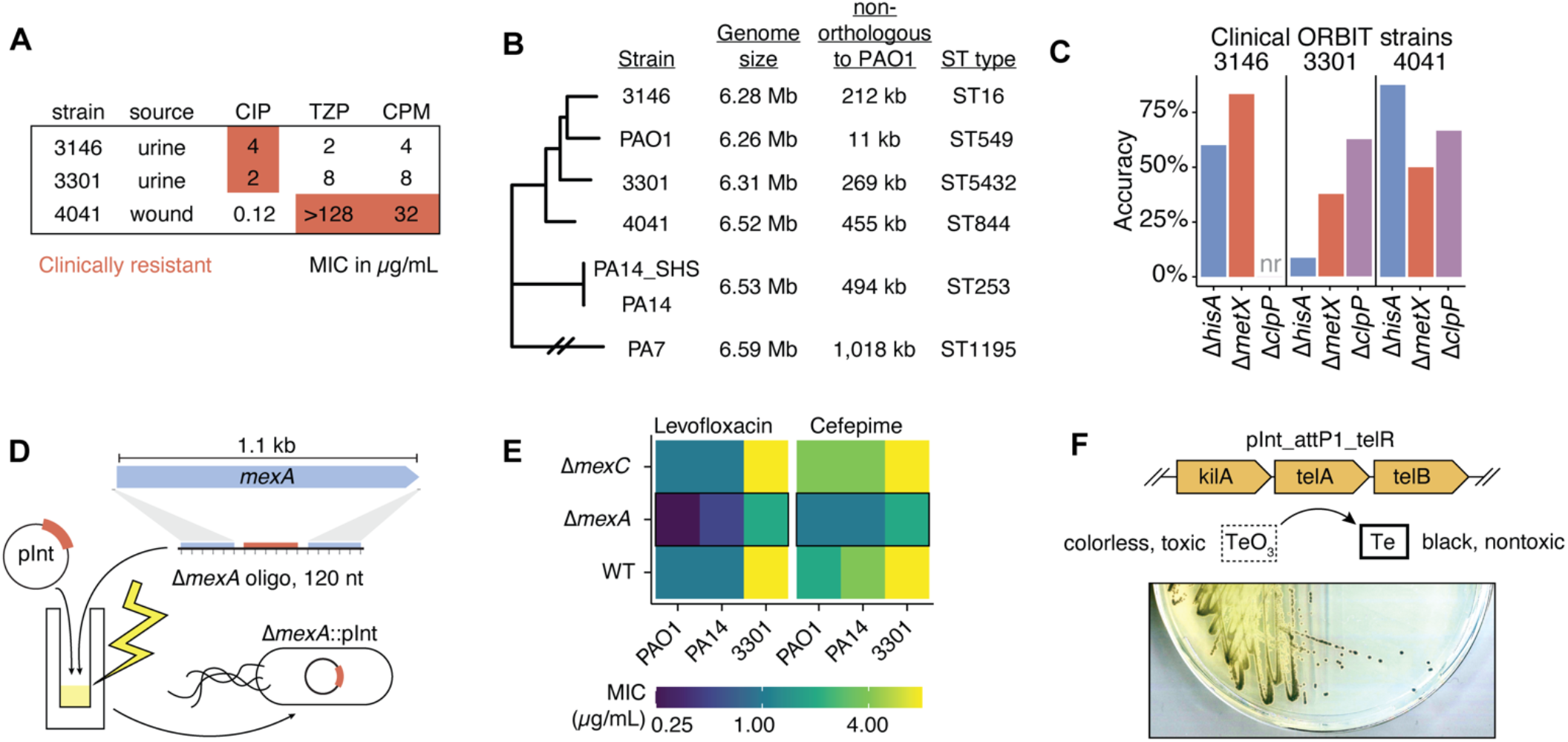
ORBIT in Pa clinical strains. A) Texas Infectious Diseases Biorepository (TIDB) clinical strains with minimum inhibitory concentrations (MIC, µg/mL) for ciprofloxacin (CIP), piperacillin-tazobactam (TZP), and cefepime (CPM). B) Whole genome phylogenetic tree for clinical isolates and lab PA strains, annotated with genome size, genomic content not orthologous to PAO1 (i.e. inferred accessory genome), and predicted ST type. PCR based accuracy for ORBIT gene deletions in each clinical strain. One strain that was not recovered is labeled “nr.” D) Diagram of efflux pump adaptor protein deletion. E) Levofloxacin and cefepime MIC values for wildtype, Δ*mexA* and Δ*mexC* in PA14, PAO1 and TIDB3301 strain backgrounds. F) ORBIT mutant using tellurite selectable marker integrating plasmid, pInt_telR. Black color shows resistant colonies producing elemental Te, converted from colorless TeO_3_.

Next, we attempted to delete the previously benchmarked test genes in these clinical isolates. Successful gene deletions were confirmed by junctional PCR for *hisA, metX*, and *clpP* loci (Fig. 3C), with accuracy ranging from 8 to 87%. Of the 9 desired strains, 8 were obtained on a single attempt (Δ*clpP* in 3146 was not observed).

To demonstrate our clinical genetic capabilities, we next decided to inactivate three of the major RND efflux pumps produced by Pa, MexAB-OprM, MexCD-OprJ and MexXY, which are known to act on various clinical drugs to various degrees (33–35). Each locus is comprised of a required adaptor protein, transporter and outer membrane channel, and we specifically deleted the adaptor protein in each locus, *mexA, mexC* and *mexX* (Fig. 3D). Including PA14 and PAO1 backgrounds, we rapidly obtained 10 of the 15 desired deletion strains and then performed antimicrobial susceptibility testing with wildtype, Δ*mexA* and Δ*mexC* for strain 3301 and lab strains. Consistent with literature that identifies MexAB-OprM as a dominant constitutive pump, only Δ*mexA* showed obvious changes in minimum inhibitory concentrations (MIC) with levofloxacin and cefepime. Interestingly, the relative fold change effects of Δ*mexA* on MIC were quite consistent across strains and drugs (2-4x), despite differences in wildtype MIC (Fig. 3E).

We also sought non-antibiotic selection markers to enable modifications in highly resistant strains. We adapted a tellurite selectable marker for use on pInt (i.e. pInt_telR), which we successfully used in PA14 (Fig. 3F). Further improvements in helper plasmid transformation and ORBIT efficiency (e.g. PMBN treatment) will be required to realize the potential of rapid pan-strain ORBIT. Nevertheless, these trials with clinical Pa strains demonstrate how ORBIT can be used in different host backgrounds to rapidly explore strain level conservation and phenotypic differences.

### An ORBIT degron system for essential hypomorph generation

To demonstrate how the Pa ORBIT system can be customized to address relevant genomic questions, we designed a degron toolkit with the goal of making essential gene hypomorphs. Hypomorphs are weakened alleles (i.e. knockdowns) that are useful for quantifying the contribution of a particular gene to both basic biology and drug resistance phenotypes (18, 36). Here, we used the native ssrA system, where the endogenous protease, ClpXP, recognizes a short peptide sequence and degrades the attached protein (37, 38). The terminal amino acids of the ssrA tag are known to modulate the level of proteolysis, so varying tag sequence can achieve gradated levels of gene knockdown. To establish the degree of ssrA mediated knockdown in strain PA14, we first tested common variants with a replicating plasmid CFP reporter (39). We observed relatively consistent knockdown levels across different copy numbers of CFP (i.e. inducer levels), and a distinct ordering of tag strength ASV<LAA<AAV<LVA (Fig. 4A). We also confirmed that degron effects were mediated almost entirely by ClpP, since no knockdown was observed in a Δ*clpP* knockout. Next, we adapted ORBIT integrating plasmids to carry these degrons, such that fusion proteins could be made by targeting the C-terminus of protein coding sequences, with in frame translation proceeding through the att linker and into the degron (Fig. 4B). We also created a “stop” control, which includes a stop codon downstream of the linker, allowing for inference of the tag-specific effect. We validated these pInt-based degrons by targeting *hisC1* (histidine biosynthesis) and found that ASV tags did not eliminate growth on minimal media, whereas the stronger tags (LAA, LVA) showed the same phenotype as a *hisC1* knockout (Fig. 4C).

**Figure 4.**
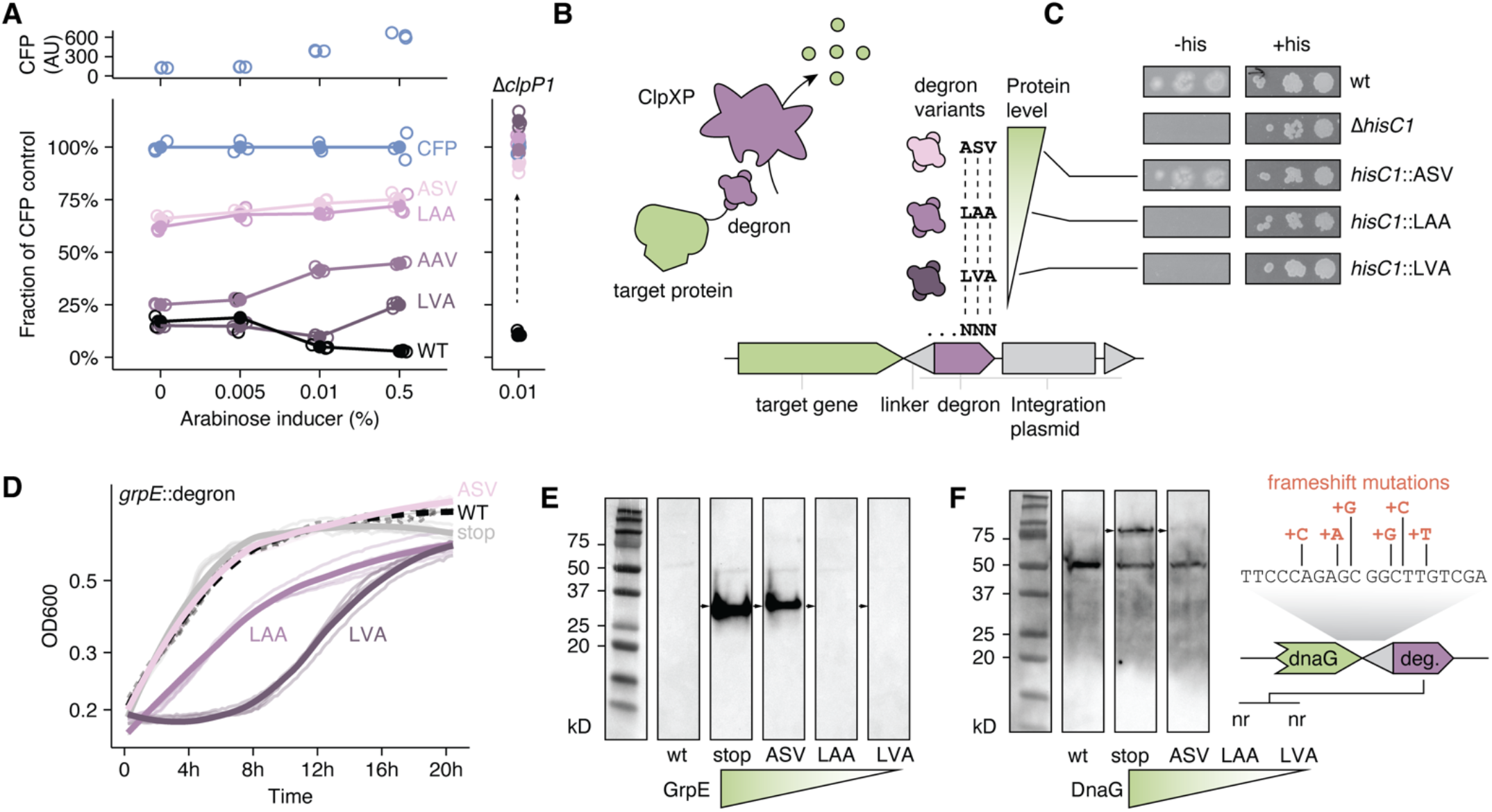
Degron toolkit for essential gene knockdowns. A) plasmid borne CFP with ssrA tag variants, ASV, LAA, AAV, and LVA showing different steady state levels of fluorescence relative to the untagged CFP control (n=3). Fluorescence was assayed at four different inducer concentrations that yielded increasing levels of CFP. Raw fluorescence of untagged CFP is shown on top in blue. All plasmids were assayed in a Δ*clpP1* background, which showed no knockdown (right). B) An ORBIT pInt-based degron system with ssrA variants. C) ORBIT degrons were tested with hisC1, which yielded variable growth on minimal medium with and without histidine. B) Essential gene *grpE* tagged with each degron or the stop control shows stepwise growth defects in LB medium. C) *grpE* tagged with each degron shows stepwise decrease in protein levels by anti-FLAG western blot. F) *dnaG* tagged with ASV shows reduced protein levels. Strong degrons LAA and LVA were not recovered during ORBIT construction. Each colony acquired a base insertion near the genome-tag junction, creating a frameshift that breaks the tag.

Next, we evaluated if these tools could be used to precisely target essential genes. We attempted to construct various degron mutants for GrpE (heat shock protein), FolA (folate biosynthesis), and DnaG (DNA primase). To evaluate the level of knockdown for each target, we also added an internal 3x FLAG tag between the pInt att site (linker) and the degron, which enabled Western blot detection. For *grpE*, we successfully recovered all degron mutants. GrpE-degron strains showed stepwise fitness effects in LB, where GrpE-ASV showed wildtype fitness, but GrpE-LAA and GrpE-LVA displayed increasingly profound growth defects (Fig. 4D). This effect was consistent with the observed level of GrpE knockdown, where ASV reduces protein levels, but LAA and LVA tagging yield undetectable levels of GrpE (Fig. 4E). For *folA*, we also obtained all four knockdown variants, however, we were surprised they did not show growth defects in LB medium (Fig. S6). We confirmed the identity of FolA-LVA with whole genome sequencing and observed protein level knockdowns by Western blot. To increase selection on FolA levels, we added trimethoprim, an antibiotic which inhibits the FolA enzyme. Under weak trimethoprim selection, the *folA* strains did exhibit stepwise growth defects, further confirming the successful modulation of essential protein levels.

For DnaG, only the stop and weak ASV tag were obtained, which showed a mild knockdown without a strong growth phenotype (Fig. 4F). The stronger LAA and LVA degron strains could not be obtained for DnaG, likely because steady state protein levels were too low to support the essential DnaG function. Few putative recombinants were obtained, and colonies that appeared correct by PCR showed a range of single base insertions near the *dnaG*-pInt junction that eliminated the ssrA tag via frameshift (Fig. 4F). Although these mutations rendered the strains unusable, we reasoned that similar mutations could be easily detected and discarded in a pooled sequencing experiment. The quantitative level of knockdown appeared to vary across protein targets, which is expected as protein-specific properties affect ClpXP accessibility (38). For example, LAA appeared to be a weak tag with CFP, but strong with all other targets. However, we were encouraged by the reproducible ordering of our tag variants and successful knockdown of highly essential genes. The high fitness of GrpE-ASV, FolA-ASV/LAA/LVA, and DnaG-ASV indicate that certain Pa essential genes can be knocked down significantly without profound consequences, raising the possibility that Pa essential gene drug targets may display different levels of vulnerability.

### Constructing and assessing a pooled hypomorph library

To create a larger set of essential gene hypomorphs and explore essential protein vulnerability, we took a pooled mutant library approach. We developed a computational pipeline and designed targeting oligos for a set of 60 core essential genes (40). These ranged in genomic position and molecular function. We prioritized cytosolic localized proteins due to ClpXP accessibility, but we included several non-cytosolic targets to test the effectiveness of our system. For the 60 targets, we ordered 150 nt oligos as a pool, which arrived as a single tube in 3 days. We performed four transformations using the oligo pool in the exact same manner as single oligos, paired with each of the four degron integrating plasmids (stop, ASV, LAA, LVA) and obtained hundreds to thousands of colonies, with LVA yielding the lowest number (Fig. 5A).

**Figure 5.**
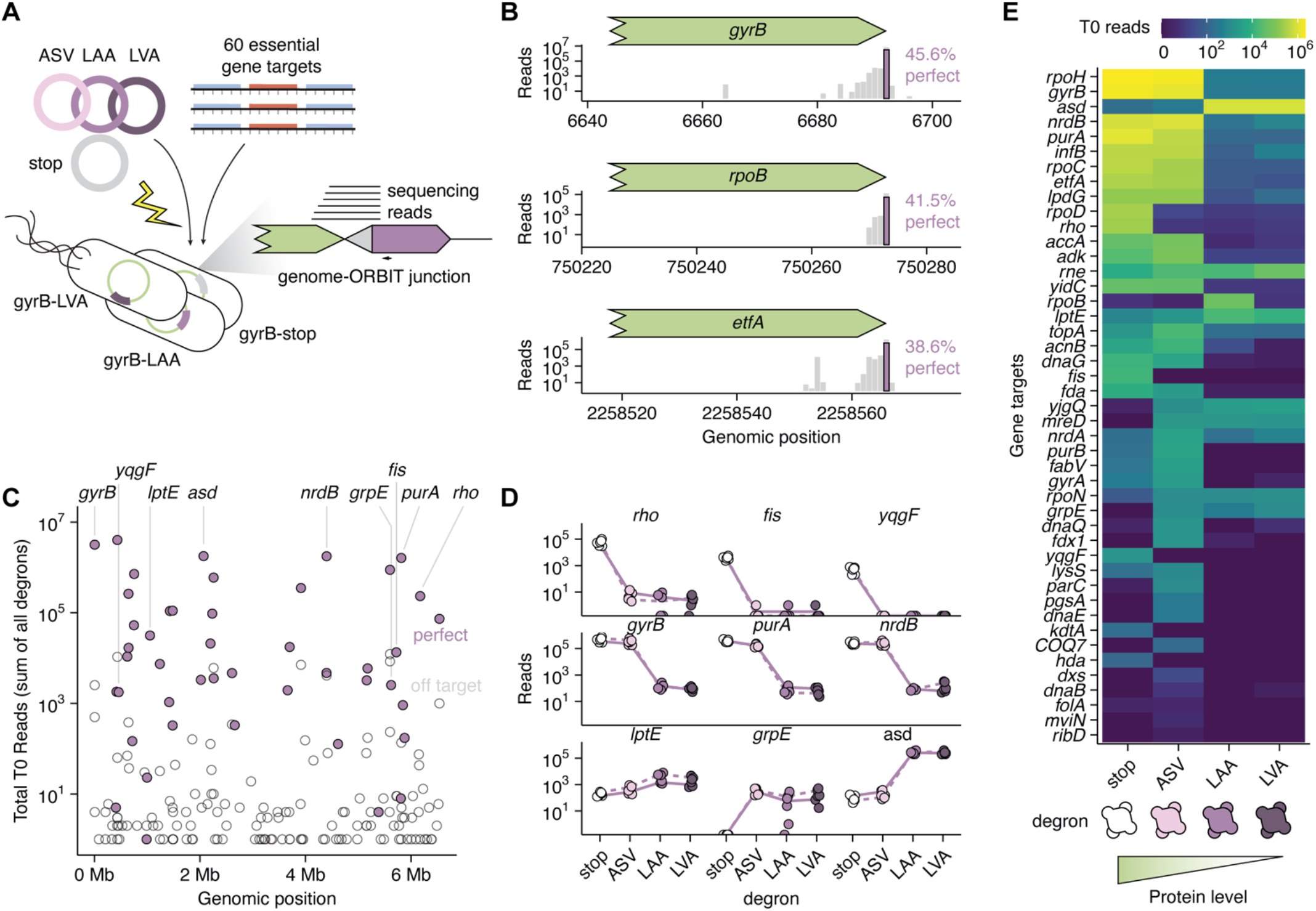
Pooled hypomorph construction and representation. A) Diagram showing the ORBIT scheme for the pooled hypomorph library. B) Representative target gene-degron fusions. Plots show reads from T0 condition pooled for all 4 degron variants and replicates. Gray bars show reads aligning to the genome with tolerance for minor errors. Purple bars show reads for all degrons observed at the correct base, with perfect adapter and genomic sequences. The perfect rate refers to the strictly perfect reads compared to all the reads shown in gray bars. C) Pooled perfect reads for all degrons and replicates for each genomic target at T0. Open circles represent off target reads that contain perfect adapters but do not match expected genomic sequences. D) T0 (dashed line) and LB (solid line) reads for gene targets across degron variants, showing vulnerable and invulnerable examples (n=3). Raw reads are shown to observe order of magnitude effects in initial mutant representation. E) Heatmap of T0 pooled perfect reads summed across replicates for each gene target-degron combination. Note the darkest blue corresponds to a read count of 0, but the color map is on a log scale (pseudocount = 1).

To evaluate the representation of strains in our pooled library, we optimized a short-read sequencing library preparation, adapted from Tn-Seq, that amplifies from a common region of the integrating plasmid and a randomly ligated adapter in the genomic DNA (16, 41, 42). We separately sequenced the stop, ASV, LAA, and LVA mutant pools using a small-scale sequencing kit. To identify and count mutants, we used the degron-linker sequence as a common handle and mapped putative genomic regions to the PA14 reference genome. Matches based on genomic position were counted as successful degron tagging events and we observed degron fusions for 39 of the 60 essential targets. We observed higher representation of stop and ASV strains compared to LAA, LVA further supporting the relative ordering of tag strength that we had previously observed.

Then we pooled all four mutant sets into a single pool (1:1:1:1 by optical density) and quantified the fitness of the mutants in LB, and sub-MIC concentrations of three clinically relevant drugs. We prepared biological triplicate libraries and performed a larger scale sequencing run that included resequencing of our initial four libraries, which let us assess our ability to differentiate degron tags based on single SNPs. This analysis demonstrated a highly reliable pipeline for identifying degron sequences, even with single nucleotide differences between LAA and LVA tags (>99.9 % accuracy)(Fig. S7A-B). High accuracy for individual targets was also observed, as exemplified by *gyrB, rpoB* and *etfA* (>96% on target, 38–45% perfect reads) (Fig. 5B). Given our difficulty obtaining DnaG degrons without mutations, we chose to only accept perfect reads that contained the exact pInt sequence, and exact genomic sequence with the junction at the exact correct base. With that high standard, a significant fraction of reads was discarded, but this process yielded a dataset that unambiguously tracks intact hypomorph strains. At all timepoints and conditions, degrons and replicates yielded highly reproducible read counts (R^2^ = 0.768 to 0.988) (Fig. S7C-D). The higher sequencing depth let us gain confident reads for 43 of the 60 targets (Fig. 5C). The 17 unobserved targets may also be of interest, since it is likely that they are very susceptible to any modification, even the linker encoded in the stop control.

To understand the baseline fitness of our hypomorphs, we examined the results of the screen for each target at the initial timepoint (T0) and LB condition (Fig. 5D-E, S8). Interestingly, we observed a wide range of degron phenotypes across target genes. Generally, we found both completely insensitive and highly sensitive targets, as well as several cases of increasing (as opposed to decreasing) read counts with tag strength. For example, the genes *gyrB* (gyrase), *purA* (purine biosynthesis), and *nrdB* (nucleotide biosynthesis) all show monotonic decreasing relationships of mutant representation and tag strength, where the weakest ASV tag does not reduce abundance, but stronger tags do. Several targets show hypersensitivity, for example, *rho* (transcription termination), *fis* (transcriptional regulator / nucleoid factor) and *yqgF* (ribosomal RNA processing) all display dramatically reduced representation even at the weakest tag strength (ASV), indicating they are highly vulnerable targets. LptE (lipopolysaccharide) and Rne (RNase) do not display any sensitivity to the tags, which could indicate they are not suitable substrates for ClpXP. This is possibly due to the secreted or membrane bound nature of LptE and Rne, however, we observe degron sensitivity in other membrane associated proteins like YidC (membrane insertase, inner membrane). Lastly, several targets show complex increasing relationships like *grpE, asd* (aspartate dehydrogenase) and *mreD* (cell division). It is likely that some of these relationships are explained by the linker itself (present in “stop” strains) causing large fitness defects, and reduction in the copy number of the defective protein improves fitness. These results show that ORBIT enables the rapid creation of pooled hypomorph libraries, and that Pa essential genes likely vary in knockdown vulnerability.

### High throughput hypomorph screening reveals synergistic targets

To investigate fitness interactions between hypomorphs and commonly used Pa clinical drugs, we next examined the pooled screen results in the presence of three antibiotics (Fig. 6A): Levofloxacin, a fluoroquinolone inhibitor of DNA replication (sub-MIC = 0.125 µg/mL); trimethoprim-sulfamethoxazole, a combination of folate metabolism inhibitors (sub-MIC = 5.6 µg/mL), and tobramycin, an aminoglycoside that inhibits translation (sub-MIC = 0.4 µg/mL)(Fig. S8). We reasoned that synergistic knockdown-drug interactions would manifest as lower fitness, while antagonistic interactions would yield higher hypomorph fitness than without the drug. Using differential abundance inference and a false discovery rate of 10%, we observed a range of significant synergistic and antagonistic relationships (Fig. 6B, S9). Many significant hypomorph-drug interactions involved levofloxacin, likely because the effective dose appeared stronger than for the other two drugs. While levofloxacin interactions may have been expected for essential proteins involved in DNA replication (e.g. *gyrAB*) or nucleotide biosynthesis (e.g. *purA, nrdAB*), we did not anticipate this result for seemingly unrelated targets (e.g. *lptE, etfA, infB*). Many targets showed degron dependent interaction. For example, LptE-LVA shows a profound defect in the presence of levofloxacin, while the other degron tags show weaker or no effect. GyrB is a known fluoroquinolone target, and *gyrB* hypomorphs displayed a complex relationship with the drug, where stop and ASV showed antagonism, LAA and LVA showed weak synergy.

**Figure 6.**
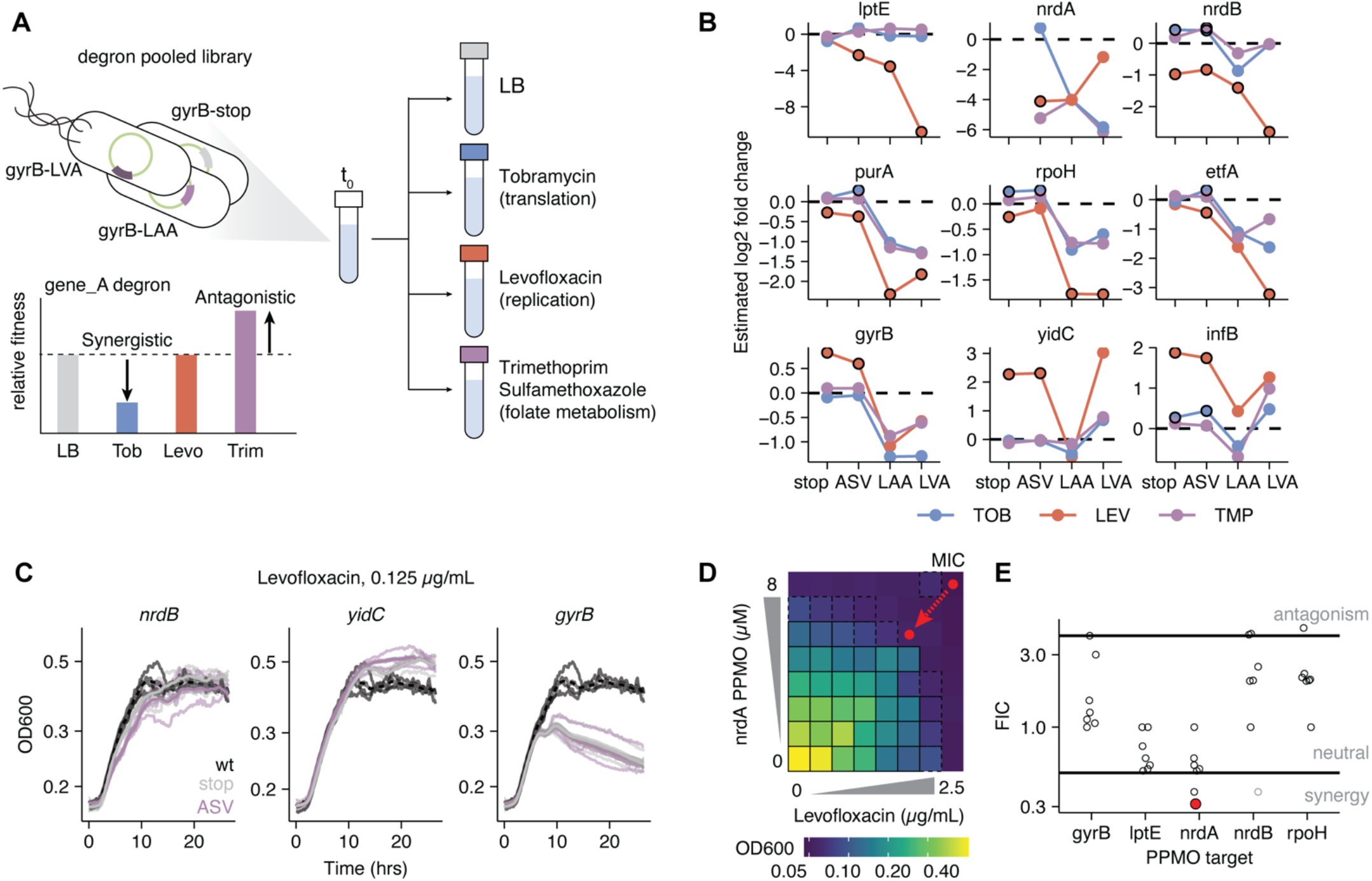
Hypomorph-clinical drug synergy. A) Diagram showing pooled screening approach and synergistic vs. antagonistic fitness interactions. B) Estimated log2 fold changes for nine gene targets across degrons (n=3). Overlaid black circles indicate significant effects (FDR<0.1). C) Growth results for individually constructed hypomorphs in levofloxacin (n=4). D-E) PPMO-levofloxacin checkerboard assays showing synergy and antagonism. FIC is fractional inhibitory concentration. PPMO and levofloxacin are diluted two-fold in adjacent rows and columns, with zero being the final concentration.

To further investigate these findings, we attempted to individually construct hypomorphs for seven additional loci: *fis, lptE, nrdB, etfA, yidC, gyrB*, and *purA*. We constructed strains with the FLAG tagged degron variants to facilitate phenotypic and western blot validation. Including the original three loci (*folA, dnaG, grpE*), we successfully made 21 of the desired 40 strains after checking a maximum of 8 colonies each. The strong LAA and LVA tags were rarely obtained, no *lptE* strains were recovered, and only the Fis-stop strain was verified. Upon Western blotting, most of these targets showed lower protein levels with the ASV tag (Fig. S6D). We reasoned that many hypomorphs may have shown interactions with levofloxacin due to differences in growth rate that may cause differences in drug susceptibility. However, we did not observe large growth rate defects in LB across the constructed strains (Fig. S6E). In sub-MIC levels of levofloxacin, we observed a mild growth rate defect for NrdB-ASV, which matches the synergy observed in the pooled experiment (Fig. 6C). YidC-ASV reached a higher optical density than wildtype, somewhat consistent with the observed antagonism in pooled format. GyrB-ASV reached a lower final optical density than wildtype, however, it shows no defect during early exponential growth in levofloxacin and a small defect in LB, consistent with the expected antagonism. Therefore, several significant hypomorph-drug interactions were confirmed with individual strains, and further work will be needed to understand the mechanistic basis of these effects.

To test if the observed hypomorph-levofloxacin interactions are relevant to co-therapy strategies, we used previously designed inhibitory antisense oligos (PPMOs) against several gene targets. PPMOs inhibit translation by blocking the ribosome binding site in a sequence specific fashion, resulting in lower steady state protein levels, like degron tags. Our hypomorph screen showed that *nrdA* (ribonucleotide reductase) was generally a very sensitive knockdown target with few reads and likely displayed some synergy with each of the tested drugs, while its molecular partner *nrdB* was less sensitive and showed strong synergy with levofloxacin. In a checkerboard assay that titrated levofloxacin and a PPMO against *nrdA* (in combination), we observed significant synergy where the addition of 2 µM PPMO drove a reduction in levofloxacin MIC from 2.5 to 0.625 µg/mL (Fig. 6D). A *nrdB* PPMO did not show significant synergy, although it is possible that strong NrdB inhibition was not achieved. Conversely, an *infB* targeted PPMO showed notable antagonism with levofloxacin, as predicted from hypomorph data, driving an MIC increase from 2.5 to 10 µg/mL (with 8 µM PPMO) (Fig. S10). LptE knockdown showed some negative interaction but was not strong enough to be classified as synergy by fractional inhibitory concentration (Fig. 6E). Other PPMO targets did not show strong interactions (e.g. RpoH). These results suggest that pooled knockdowns accurately predict some PPMO inhibition results, but further work is likely needed to quantify knockdown levels to make more reliable comparisons.

Collectively, these results establish degron tagging as a robust, tunable approach for probing essential gene function and vulnerability in Pa. The complete results from our hypomorph screen are included as supplementary data, and the pooled degron library is available upon request, which we hope will be a useful community resource. The high throughput ORBIT approach taken here should be broadly applicable to genome scale problems in this important pathogen.

## Discussion

In this work, we adapted an oligo based genetic toolkit to perform low and high throughput genome modifications in the opportunistic pathogen, *Pseudomonas aeruginosa*. At high throughput, we used ORBIT with degron integrating plasmids to create essential gene hypomorphs, successfully constructing 166 pooled strains with four transformations. The results of this degron screen show that Pa essential genes have variable knockdown vulnerability. For example, some target strains dropped in abundance by orders of magnitude in the presence of even the weakest, ASV, tag. These hypersensitive targets included known drug targets like *rpoD* (housekeeping sigma factor) and *rho*, but also potentially novel targets, *fis* and *yqgF*. It is interesting that the nucleoid associated protein, Fis, is sensitive to knockdown, because it is known to vary widely in copy number and is dependent on growth phase (43). Transposon screens and our work identify Fis as a highly essential protein in Pa, while it is completely dispensable in *E. coli*. Here we establish that native Fis can be 3x FLAG tagged (i.e. verified “stop” strain), which should facilitate further study of this important, yet understudied Pa protein.

We also observed significant hypomorph interactions with clinical drugs, which could inform co-therapy strategies. Notably, inhibition of many gene products reduced the effectiveness of levofloxacin, underscoring the complexity of combination therapy and underlying Pa essential gene networks. However, we also observe substantial synergy, including a potentially useful combination therapy of a *nrdA* PPMO and levofloxacin. Given this success, we believe further work should be done with the full range of essential gene targets to characterize target vulnerability and possible co-therapeutic strategies.

Future work should consider some of the shortcomings of our degron approach. First, our degron tags resulted in constitutive knockdown, which is likely the reason many strong degron strains could not be obtained. This also resulted in lower abundance of strong knockdowns, which limited our ability to detect drug interactions for very sensitive targets. One solution could be to implement an inducible degron system, with either the protease or an adaptor protein being modulated (36, 44). Importantly, our constitutive tag-based gradation approach enabled pooling of different strength knockdowns and therefore a simple differential abundance inference, but new inducible systems could be designed that preserve both benefits. Our antibiotic fitness results also demonstrate the importance of specific conditions – levofloxacin likely showed the most interactions because it was used at the highest effective dose. Future work should take advantage of the compact size of ORBIT libraries (since they are precisely targeted) to capture fitness effects at a range of drug concentrations and timepoints. These assays should also be compatible with various extensions, like animal models, persister/tolerance time kill assays, and Sort-Seq with physiological reporters (3).

Some work has been done with CRISPRi in Pa to perform knockdown screens, which we see as valuable and orthogonal to our degron approach (6–8). Both techniques offer a way to create targeted knockdowns, but they occur at different molecular levels (transcriptional or post-translational). CRISPRi guide RNAs function in trans, while degron tags necessarily modify endogenous essential gene loci, which can limit targets due to polar effects, fusion protein defects, or protease accessibility. Future degron work could limit polar effects by encoding degrons on the targeting oligo and excising the pInt. By counting perfect degron reads, our assay has some amount of verification built in that the tag is truly attached to the target protein. In contrast, CRISPRi screens rely on sequencing guide RNAs as a proxy for knockdown strain abundance. Because it is not definitively known which guide RNAs work most effectively for each target gene, or how prevalent off target effects are, most screens use multiple guide RNAs per gene target and rely on consensus, greatly expanding the size of mutant libraries. Further, CRISPRi can be silently inactivated in certain cases, skewing inferred knockdown effects (45). Therefore, both techniques offer advantages and disadvantages, and we strongly support the continued development and application of both these powerful tools. Importantly, beyond knockdown screens, ORBIT can be used to make countless types of targeted genomic modifications and be flexibly applied to many other questions.

At low throughput, ORBIT has proven very reliable to construct individual mutant strains in Pa – throughout this work we successfully constructed 77 individual strains. Here, we establish a plasmid toolkit and protocol that should enable many labs to adopt this method for routine genetics using commercially available DNA oligos. Further improvements to Pa ORBIT should include a more streamlined scarless modification strategy, characterization of orthogonal att sites for sequential modification, improved efficiency (particularly for clinical strains) via PMBN or other variables, and reduced targeting oligo concentration. While there are countless potential enhancements, it seems clear from our personal communications with members of the Pa research community that the ORBIT tools described here will enable new biological questions and genomic studies. We believe ORBIT offers an optimal strategy to create arrayed mutant libraries (e.g. like *E. coli* Keio collection) and sophisticated pooled mutant libraries, resources that could impact Pa research broadly.

Beyond a single species of bacteria, ORBIT now offers a nearly identical genetic logic and implementation that can be seamlessly applied at low and high throughput across multiple bacteria. As we hope the ORBIT ecosystem continues to grow, we expect that this inherent compatibility will only become more powerful as our group and others extend and modify the ORBIT components to function in new organisms and address new biological questions.

## Materials and methods

### Routine culture, cloning, and transformation

For routine culture, Pa and *E. coli* strains were inoculated directly from glycerol stocks into 3 mL of LB medium and typically cultured at 37°C in shaking incubators overnight (225 RPM, Innova 42). When required, minimal medium was M9 with either glucose or succinate as the carbon source and amino acid supplementation (histidine or methionine) was added to 40 µg/mL. As appropriate, liquid cultures and agar plates were supplemented with 300 µg/mL carbenicillin (100 µg/mL for *E. coli*), 80 µg/mL gentamicin (15 µg/mL for *E. coli*), 900 µg/mL kanamycin (35 µg/mL *E. coli*), or 30 µg/mL potassium tellurite (6 µg/mL for *E. coli*). Molecular cloning of plasmids was performed using the NEB HiFi DNA assembly kit and standard replicating plasmids were maintained in *E. coli* DH5a, while ORBIT integrating plasmids were maintained in a pir+ host strain (BW25141). Helper plasmids were typically prepared via standard miniprep, while larger quantities of integrating plasmid were prepared with the Zymo MidiPrep kit using the low copy plasmid protocol.

To introduce the ORBIT helper plasmid into Pa strains, a standard electroporation was performed. Cells from an overnight culture were diluted and grown to exponential phase (OD 0.5), then they were washed three times in sterile 1 mM MgSO_4_. Approximately 100 ng of purified plasmid DNA was used, and electroporation was performed with 2 mm gap cuvettes using 2.5 kV (25 μF, 600 Ω, exponential decay waveform). Cells recovered for 2 hours (6 hours for clinical strains) in 1 mL of SOC medium shaking at 37°C and then plated on selective medium, typically gentamicin for pHelper_PA_std.

### ORBIT strain construction

A complete protocol for constructing and verifying ORBIT strains is included as supplemental material. Briefly, targeting oligos were designed containing homology arms immediately adjacent to the gene to be deleted. For translational fusions, the stop codon was deleted, and adjacent homology arms were used. To target the lagging strand template, the minus strand genomic sequence was used for the oligo when the target fell within replichore 1 (i.e. origin to terminus), and the plus strand genomic sequence was used when the target fell within replichore 2 (i.e. terminus to origin). The attB sequence was always designed to face the same direction as the gene, where ‘GGCTTGTCGACGACGGCGGTCTCCGTCGTCAGGATCAT’ is considered the forward direction. To facilitate oligo design, a web tool was developed for the PA14 and PAO1 model strain genomes (https://saunders-lab.shinyapps.io/TO_design_PA14/). Typically, oligos were designed with the web tool and then checked manually in Benchling. Oligonucleotides were ordered from IDT (ultramers) and Sigma Millipore using standard synthesis with no additional purification. Oligos were resuspended to 100 µM stock concentrations in molecular grade water and used directly in ORBIT transformations. Typically, 120 nt oligos were used unmodified, but for the PO bonded oligos two modifications were added on the 5’ end and two modifications were added on the 3’ ends of the oligos. Clean deletion oligos did not include the attB site and were 82 or 150 nt in length.

Strains that already contained pHelper were prepared for ORBIT electroporation by diluting into antibiotic containing medium from overnight cultures and grown to exponential phase (OD 0.3) and PapRecT was induced by adding 1 mM m-toluic acid (Sigma #T36609) for 30 min. Then cultures were pelleted and washed three times in 1 mM MgSO_4_ at room temperature. Typically, 9 µL of 100 µM oligo stock (900 pmol) was added to 100 µL concentrated cell aliquots, along with 100-500 ng of midiprep purified integrating plasmid. Co-electroporation was performed with 2 mm gap cuvettes using 2.5 kV and cells typically recovered in 1 mL of SOC medium shaking at 37°C overnight. The next morning cells were concentrated and plated on selective media either by streak plating or drip plating to facilitate CFU counting (10 µL serial dilutions).

To cure the standard helper plasmid, colonies were restreaked on 10% sucrose (in no salt LB). Colony PCR was frequently used to verify ORBIT genotypes either by amplifying the integrating plasmid-genome junctions (i.e. junctional PCR) or by amplifying the entire ORBIT modification using genomic primers (i.e. spanning PCR). Standard PCR was performed using GoTaq Green V2 (Promega) and products were sequenced with Sanger, as needed. To detect off target SNPs in whole genomes, genomic DNA was extracted using a bacterial/fungal gDNA miniprep kit (Zymo D6005) and submitted to SeqCoast for Illumina whole genome sequencing. Raw reads were compared against a PA14 reference genome with breseq (see GitHub for script)(46).

### Clinical Pa strain ORBIT and PPMO checkerboard assays

Bacterial strains were laboratory strains or clinical isolates chosen from the Texas Infectious Diseases Biorepository (TIDB) (IRB-approved study protocol # STU-2018–0319). Ten Pa strains were manually selected from the TIDB based on their expected sensitivity to gentamicin and carbenicillin. Strains showed considerable phenotypic heterogeneity in pigment, polysaccharide production, and growth rate. Helper plasmid transformation was performed as with lab strains, plating on 80 µg/mL gentamicin. To confirm helper plasmid replication, putative strains were grown in selective gentamicin and then subject to a native miniprep using a standard kit. Plasmid extracts were run on an agarose gel and sequenced via whole plasmid nanopore sequencing (Plasmidsaurus). Only three strains grew and maintained the helper plasmid intact, TIDB 3146, 3301 and 4041. Standard ORBIT was performed for *hisA, metX* and *clpP* targets in these strains using 150 nt targeting oligos that were designed for PA14 (target strains had the same homology arm sequences). For each putative recombinant strain, 8 colonies were tested by PCR. A similar approach was taken for efflux pump genes *mexA, mexC*, and *mexX*.

To phenotype efflux pump mutants, strains were grown overnight and then resuspended in 96 well plate cultures containing two-fold serial dilutions of clinical antibiotics. After 24 hours of growth at 37°C, OD600 was read on a plate reader. MIC was determined as the minimal amount of antibiotic that prevented an OD of less than 0.06. Checkerboard assays were performed using levofloxacin and PPMOs against several targets. PPMOs were designed using a custom alignment tool as previously described and then synthesized by Sarepta Therapeutics (25). Two-fold dilutions of both compounds were performed in combination across the rows and columns of a 96 well plate. After 24 hours of growth at 37°C, OD600 was read on a plate reader. MIC was determined as the minimal amount of antibiotic or PPMO that prevented an OD of less than 0.06. FIC was calculated as FIC = (MIC_PPMO_with_levo_ / MIC_PPMO_only_) + (MIC_levo_with_PPMO_ / MIC_levo_only_).

### Clinical Pa strain genome sequencing and analysis

Genomic DNA from TIDB clinical strains of interest (3146, 3301, and 4041) was extracted using the ThermoScientific GeneJET Genomic DNA Purification Kit (#FERK0722). Sequencing libraries were prepared from 20 ng/µL gDNA aliquots using the Rapid Barcoding Kit (SQK-RBK114.24; Oxford Nanopore Technologies) according to vendor’s instructions. Tagmentation (with unique barcodes) was performed by incubating at 30°C for 2 minutes, then 80°C for 2 minutes. The barcoded samples were then pooled in equimolar amounts, cleaned using AMPure XP beads, and washed twice with 80% ethanol. The pooled library was eluted in elution buffer, and Rapid Adapter was ligated to the pooled barcoded DNA by incubation at room temperature for 5 minutes. The final library was quantified using the Qubit fluorometer (dsDNA HS assay), further mixed with sequencing buffer and library beads, then loaded onto a primed R10.4.1 flow cell (FLO-MIN114) on a MinION device (Oxford Nanopore Technologies). Sequencing was conducted using MinKNOW software for 72 hours.

Nanopore signal data were basecalled with Dorado using the HAC model and the corresponding sequencing kit option. Basecalled reads were demultiplexed with Dorado, and barcode-specific BAM files were converted to FASTQ. Reads shorter than 1 kb were removed, and the remaining reads were filtered with Filtlong, retaining the top 90% by read quality. Filtered reads were assembled with Flye, followed by one round of Racon polishing using minimap2 read-to-assembly alignments and final polishing with Medaka. Assembly quality and contig lengths were summarized with QUAST and custom sequence-length scripts. Taxonomic composition was assessed using Kraken2 with the standard database, and multilocus sequence typing was performed with mapMLST. Coding sequences were predicted with Prokka. Filtered reads were aligned back to the polished assemblies with minimap2, and the resulting sorted BAM files were used to estimate assembly depth. Additional antimicrobial-resistance-related analyses were performed by translated read mapping against an ARDB/CARD-derived protein database using DIAMOND BLASTX and custom ORF/protein mapping scripts (47). A whole genome phylogenetic tree including clinical and lab strains was constructed with parsnp using fasta genome files (48). FastANI was used to compare each genome to PAO1, and the non-orthologous genomic fraction was inferred by subtracting the total orthologous genomic fragments from the total genome length for each strain (49).

### Degron strain construction and validation

Integrating plasmids (pInt_attP1_ampR backbone) were modified to carry ssrA tags immediately downstream of the attP site, such that translation proceeding through the att linker would result in an in-frame tag. Therefore, targeting oligos for degron tagging were designed to integrate the attB site in frame with each gene coding sequence, inserting constructs immediately before the stop codon. Versions of the integrating plasmids were created that also carry 3x FLAG epitopes upstream of the degron tag, enabling Western blot detection. To prepare degron strains for Western blotting, they were inoculated overnight in LB + carbenicillin and then diluted 1:1000 into LB before pelleting at OD ∼1.0. Pellets were resuspended in 100 μL of Laemmli Buffer + 5% fresh Beta-mercaptoethanol and boiled in microfuge tubes for 20 min. Any kD mini protean TGX 10 well gels were used from Bio-Rad (#4569033). Gels were run in 1x SDS PAGE buffer with Precision protein plus ladder (Bio-Rad). Typically, 25 μL of sample was loaded into each well. Gels were run for 30-40 min at 200V and then transferred to Nitrocellulose membranes. Membranes were dried for 15 min, then blocked with 10 mL TBST + 3% milk on an orbital shaker overnight at 4°C. An HRP conjugated primary anti-FLAG antibody (1:100) (Abcam #ab49763) was used to probe membranes by incubating 1 hour at room temperature on the orbital shaker. After washing 3x with TBST (15 min each at room temperature), chemiluminescent reagent was added (SuperSignal West Atto – Thermo Scientific #A38554). Membranes were placed in a sheet protector and imaged on a Bio-Rad ChemiDoc system.

The 60 essential gene targeting oligo pool was ordered as an oligo pool (50 pmol/oligo), where each 150 nt oligo was designed computationally following the rules mentioned above. Pooled degron construct was performed in the same manner as individual ORBIT modifications, where the oligo pool was resuspended to 100 µM and used directly in transformations with the four different degron integrating plasmids. After spread plating, colonies were resuspended off agar plates, pooled, and saved as glycerol stocks. Cell pellets of each pool were also directly saved for sequencing validation. Pooled fitness assays started by inoculating aliquots of the stop, ASV, LAA, and LVA pooled glycerol stocks overnight in carbenicillin. The next morning subpool cultures were measured for OD and pooled 1:1:1:1 to a final OD of 0.1 in three separate flasks (biological replicates). These 50 mL carbenicillin cultures were grown to OD 1.0 and served as the T0 samples. Then, each replicate was separately diluted into 50 mL LB only, LB + levofloxacin (0.125 µg/mL), LB + trimethoprim-sulfamethoxazole (5.6 µg/mL), or LB + tobramycin 0.4 µg/mL) to an OD of 0.01. Cultures were grown to an OD of 1.0, which took longer to achieve for levofloxacin. Therefore, we inferred that levofloxacin was used at a higher effective dose than other antibiotics, despite our efforts to choose similar dosages that yielded only mild growth defects.

### Hypomorph pooled sequencing and analysis

Library preparation for ORBIT mutant pools was performed in a manner analogous to Tn-Seq. Genomic DNA (gDNA) was extracted from mutant library cell pellets using a bacterial/fungal gDNA miniprep kit (Zymo D6005). DNA was fragmented and adapters were ligated using the NEB Ultra II FS Library Prep Kit for Illumina (#E6177). Fragmentation was performed for 10 min at 37°C and magnetic bead size selection was performed for fragments sized 275 to 475 bp. PCR1 was performed with 500 ng of adapter ligated DNA using a primer that binds the integrating plasmid and a primer (with overhang) that binds the adapter using Q5 ultra polymerase (98°C – 10 s, 68°C – 30 s, 72°C – 45 s, 13 cycles). Illumina compatible overhangs were added with PCR2 using 10 ng of the purified PCR1 product (98°C – 10 s, 65°C – 90 s, 14 cycles). Products were cleaned with magnetic beads after each PCR and served as the final sequencing library. Libraries were quantified with Qubit fluorometry (Thermofisher # Q32851) and typically pooled 1:1. Initial sequencing libraries were run on an Illumina MiSeq instrument with a Nano V2 kit (300 cycles total, 200 cycles Read 1). A larger subsequent pooled sequencing experiment was sent to Azenta and run on an Illumina NovaSeq instrument.

First, demultiplexed sequencing reads were quality filtered using default settings and fastp. Then ORBIT mutations were identified by treating them as adapters in Cutadapt. Each degron sequence (stop, ASV, LAA, LVA) was treated as a separate adapter and Cutadapt categorized reads into separate degron files. Perfect reads contained no errors or mismatches in this adapter trimming step, while initial “imperfect” reads tolerated 2 base errors. Following trimming of the integrating plasmid sequences, putative genomic sequences were mapped to the PA14 genome. Then genomic positions were matched between mapped reads and designed constructs to count the number of reads observed for each targeted strain. Perfect reads were identified by filtering for paired read alignments where read 1 contained no mismatches or indels. Sequencing reads were processed on the BioHPC on a node with 128 GB of RAM and 32 logic cores. The GitHub repository contains the sequencing read processing scripts.

To obtain fitness values for each mutant across clinical drug conditions, differential relative abundances were inferred with edgeR (50), treating each gene target-degron variant as a separate entity. Clinical drug abundances were contrasted with LB abundances using glmQLFTest.

## Data availability

Read counts and estimated fold changes for the pooled hypomorph libraries are included as supplementary data. Short read sequencing data from pooled hypomorph libraries will be uploaded to the Short Read Archive, as will the long-read data used to sequence the TIDB clinical strain genomes. Raw data used to construct the figures (e.g. colony counts) are included in the GitHub repository. R scripts used to generate the figures from the data are also included in the code repository (https://github.com/saunders-lab/pa_orbit). Any other data is available from the authors upon request.

## Acknowledgments

We acknowledge internal funding to SHS from the Lyda Hill Department of Bioinformatics and Green Center for Systems Biology (University of Texas Southwestern Medical Center), as well as funding to DEG (Kleberg Family Foundation and NIH AI41101). We thank Kimberly Reynolds for feedback on the manuscript and access to a variety of laboratory equipment. We thank David Basta for performing initial Pa ORBIT assays, and early support from Dianne Newman and the Center for Environment Microbe Interactions (California Institute of Technology).

