## Supporting information for "Oligonucleotide genetics for *Pseudomonas aeruginosa* enables high throughput hypomorph screening"

**This PDF file includes:**

Supporting text. *Pseudomonas aeruginosa* ORBIT protocol
Figure S1. ORBIT Efficiency details
Figure S2. Gene deletion benchmark details
Figure S3. Sanger sequencing of ORBIT modified loci
Figure S4. Markerless mutations and alternative att sites
Figure S5. Alternative helper plasmids and large integrations
Figure S6. Individual hypomorph phenotypes
Figure S7. Degron identifiability and reproducibility in pooled sequencing assays
Figure S8. Degron read count overview
Figure S9. Degron fitness effects across conditions
Figure S10. PPMO checkerboard assays
Table S1. Plasmids used in this work
Table S2. Strains used in this work

**Other supporting materials for this manuscript include the following:**

Table S3. Oligonucleotides used in this work
Dataset S1. Pooled hypomorph targets, read counts and fold changes

### ***Pseudomonas aeruginosa* ORBIT protocol**

#### Electrocompetent cells with induced helper plasmid + ORBIT integration

This protocol provides simple instructions for preparing induced electrocompetent cells and using them for ORBIT modifications. It is assumed that you already have your strain of interest with the ORBIT pHelper plasmid. It is also assumed that you already have an appropriate targeting oligo and integrating plasmid. See the Saunders Lab website for additional resources on ORBIT.

**Note: This entire procedure is performed at room temperature (no ice). To our knowledge, the electrocompetent cells cannot be stored in the -80, they must be used immediately. These are important differences compared to *E. coli*.**

This procedure was written for a 50 mL culture, enough for 4x transformations, but it can be scaled up or down to whatever volume is required. Also, labs make competent cells in slightly different ways. We have tested different electrocompetent cell prep protocols and we are confident in the one provided, but other variations might also work fine. Feel free to modify as necessary.

*Please use safety precautions while working with this BSL2 organisms, including PPE (lab coat, safety glasses, and gloves). All steps should be performed in the tissue culture room and the biosafety cabinet whenever possible.*

**Materials**

- Strain of interest with helper plasmid
- 1M m-toluic acid in ethanol (Sigma #T36609), inducer
- 1 mM $MgSO_4$, wash solution (sterile – typically autoclaved)
- SOC + 1% arabinose, recovery medium
    - SOC medium (per L: 5 g yeast extract, 0.5 g NaCl, 20 g tryptone, after autoclave add 20 mL of 1 M glucose)
    - 20% arabinose, inducer (filter sterilized)
- 2 mm electroporation cuvettes
- 100 µM ORBIT Targeting oligo in water
- ~100 ng / µL integrating plasmid in water (we use Zymo Midiprep low copy protocol)
- 10% sucrose plates with no salt LB (For 500 mL: autoclave 125 mL H2O + 50 g sucrose (40% wt/vol) and separately 375 mL $H_2O$ + 5 g tryptone, 2.5 g yeast extract and 7.5 g agar. Cool separate sucrose and LB solutions at 60°C and mix before pouring.)

Common materials
- LB broth
- 250 ml flasks

Common equipment
- Shaking incubator
- Electroporator
- 50 mL conical centrifuge

#### Competent cell prep

1. **Grow an overnight culture of the helper plasmid strain.** From a frozen stock or a colony, grow a 3 mL LB culture of the helper plasmid strain and make sure to supplement with the appropriate antibiotic. Gentamicin 80 µg / mL is standard for the current pHelper. If using the temperature sensitive plasmid, we recommend 40 µg/mL gent at 30°C.

*The next morning*

2. **Inoculate the main culture with the overnight (1:100).** Prepare a 250 mL flask with 50 mL of LB (or scale accordingly) supplemented with the appropriate antibiotic and add cells from the overnight culture 1:100. Grow at 37°C with shaking until the culture reaches OD600 ~0.3. This should take about 3 hours.

3. **Induce with 1 mM m-toluic acid for 30 min.** Add m-toluic acid directly from the 1M ethanol stock into the culture 1:1000 for a final concentration of 1 mM (i.e. 50 µL for a 50 mL culture). Let the cells continue to grow and induce in the shaking incubator for 30 min. This step induces oligo recombineering with recT.

4. **Spin the cells down at room temperature.** Transfer 45-50 mL of culture into 50 mL conical vials. Spin the tubes at 5,000x rcf for 5 min. Pellets should be small (compared to plasmid preps), but clearly visible along the wall of the tubes. Pour off the supernatant.

5. **Wash 3x with washing solution at room temperature.** With a serological pipette resuspend the pellet in 45-50 mL of washing buffer (1 mM $MgSO_4$). Again, spin the tubes at 5,000x rcf for 5 min, and carefully pour off the supernatant. Repeat 2 more times for a total of 3 washes.

*During the final wash steps, start setting up the microfuge tubes for cell aliquots (Electroporation step 1).*

6. **Resuspend cells.** Resuspend the pellet from each 50 mL conical in the residual washing solution and bring up to a volume of ~350 µL with wash solution. This is sufficient for 4x electroporations. Electrocompetent cells must be used promptly – they cannot be saved at -80°C to our knowledge.

#### Electroporation

1. **Aliquot cells for electroporation.** For each planned electroporation, aliquot 79 µL of the resuspended washed cells (step 6 above) into a room temperature microfuge tube.

2. **Add targeting oligo and integrating plasmid.** To each cell aliquot, add the appropriate targeting oligo and integrating plasmid. Generally, 9 µL of targeting oligo (100 µM stock) and 3 µL of integrating plasmid (~100 ng/uL) are added. We recommend "no oligo" controls whenever possible.

3. **Transfer to 2 mm cuvette and electroporate.** Gently pipette the cell + oligo + plasmid mixture several times to mix and transfer to a 2 mm electroporation cuvette. Electroporate at 2.5 kV (25 µF, 600 Ω, exponential decay waveform). Note, the cuvette and electroporation conditions are a bit different than for *E. coli*.

4. **Recover in SOC + arabinose overnight.** For each cuvette, immediately add 1 mL recovery medium following electroporation and then transfer cells + medium to culture tube for shaking at 37°C overnight. The recovery medium should be prepared in advance and used at room temperature. Make SOC + 1% arabinose by diluting 20% arabinose (sterile) into SOC medium.

*The next morning*

4. **Plate on integrating plasmid antibiotic.** For maximum recombinant yield, pellet the culture (5,000 rcf for 5 min) and resuspend in ~100 µL LB. Then plate the entire volume on the appropriate antibiotic agar plate. Typically, this is Carbenicillin 300 µg/mL for pInt_ampR. We typically observe 100-1000 colonies, depending on the targeting oligo and locus.

One convenient way to plate is to pipette the liquid at one edge of the plate and then streak the liquid using classic microbiological technique. This is guaranteed to yield single colonies regardless of the number of recombinants. On the other hand, it is very difficult to determine efficiency or compare quantitatively, so plating with beads or drip plates (our favorite) is useful. Drip plates are made by serially diluting a culture in a 96 well plate and then using a multichannel to spot 10 µL spots and tilt to form easily countable "drips."

5. **Verify mutants & cure pHelper.** These initial recombinants should have the ORBIT modification if everything went well. This can be immediately verified by performing colony PCR with short primer sets for the genome-integrating plasmid junctions and further verified with Sanger sequencing. At this stage mutants still have the helper plasmid, so correct colonies should be cultured in liquid with integrating plasmid antibiotic and then plated for single colonies on LB (no salt) + 10% sucrose. After helper curing, ORBIT modifications should be completely stable and final verification can be performed, ideally PCR + Sanger spanning the entire modified locus.

We recommend GoTaq green mastermix from Promega for colony PCR.

**FAQs:**

**How should I transform the replicating helper plasmid?**

*Replicating plasmids generally transform well. The helper can be directly transformed into cells from an overnight culture. Simply wash and electroporate using the 3x washes in MgSO4 as described in step 4-6 above. In other words:*

- *Grow PA14 strain of interest overnight in 3mL.*
- *Centrifuge culture for 5 min @ 5,000 x g (room temp).*
- *Wash 3x with 1mM MgSO4.*
- *Resuspend and electroporate with 100 ng replicating plasmid.*

**How can I confirm the helper plasmid went into my strain?**

*We like to perform a miniprep of the actual strain and run the supercoiled plasmid directly on a gel next to the E. coli miniprepped plasmid. If there is any doubt, whole plasmid sequencing can be confirmed. Otherwise, the correct antibiotic resistance can be confirmed by streaking on plates, or a colony PCR for a specific region of the helper plasmid could be performed.*

**How should I prepare the integrating plasmid?**

*The higher the purity and concentration of the integrating plasmid prep, the better. We recommend the Zymo midi prep kit, using the low copy plasmid protocol. These kits typically yield preps >100 ng/uL. It can be worth running the supercoiled plasmid on a gel to check it does not contain high levels of high molecular weight genomic DNA, but note that supercoiled plasmid runs differently than PCR products.*

**What if I'm using the temperature sensitive helper?**

*The temperature sensitive helper replicates at 30°C, therefore all growth and recovery steps should be performed at 30°C instead of 37°C. To cure this plasmid, simply grow at 42°C – there's no need for sucrose. Importantly, we find that strains are more sensitive to gent with TS plasmids, so we recommend 40 µg/mL instead of the standard 80 µg/mL for all selective growth steps.*

**What if I want to make a clean deletion?**

*To make a clean deletion, two separate oligo incorporation steps will be performed. One step will make the modification of interest (e.g. gene deletion) and integrate the pInt_sacB plasmid. In the second step, an oligo (80-150 nt) without the attB site will be used to precisely remove the pInt_sacB integration using sucrose counter selection. In our hands using the traditional pHelper, clean deletions are possible, but can be difficult to obtain. The most common product is the locus with only the 38 bp attB site remaining, caused by the low excision activity (as opposed to integration activity) of Bxb-1 coming from the helper plasmid (even though it is uninduced). Currently, we recommend using the temperature sensitive pHelper for the original ORBIT (with pInt_sacB), then remove it at the non-permissive temperature. Then introduce the pHelper with only RecT (no Bxb-1). At this stage, ORBIT competent cells can be induced, prepared, and transformed with the clean deletion oligo and plated on sucrose following recovery. There are two other good options to make markerless (scar remaining) mutants (see below).*

**How do I make a markerless deletion?**

*There are two ways to make markerless mutations that have no antibiotic marker but do have scars. First, an ORBIT modification made with pInt_sacB can be made markerless simply by selecting on sucrose. This should get rid of the plasmid and force the excision product (Bxb-1 can both integrate and excise), leaving only a 38 bp scar. However, this attB site will be active, and likely preclude any other ORBIT modifications in this strain. Another option is to make an ORBIT modification with a standard pInt, then remove pHelper, and add a FLP expressing plasmid (e.g. pFLP3). The FLP recombinase will excise everything between the FRT sites on the pInt, leaving a scar of ~170 bp. This scar should be inert for future rounds of ORBIT.*

**Can I use less oligo?**

*Yes, you can likely use half as much (4.5 uL of 100uM stock) with little loss of efficiency. Lower amounts, probably yield significant reductions in efficiency.*

**Does the oligo length matter?**

*Yes, we generally recommend oligos that are 120 nt in length (total, including 38 bp attB site). This is mainly due to cost – these oligos cost us $24 from Millipore sigma. Longer oligos, 150 nt, tend to work with higher efficiency, but these tend to be quite expensive (when ordering individually). For situations where cost is not an issue, we would recommend such oligos from IDT (Ultramers). We do not recommend shorter oligos < 120 nt, but feel free to try if you want!*

**Can I recover for a shorter interval?**

*Yes, you can likely recover for shorter intervals. In our hands, overnight recovery is the most reliable, but a couple hours are probably sufficient for some applications.*

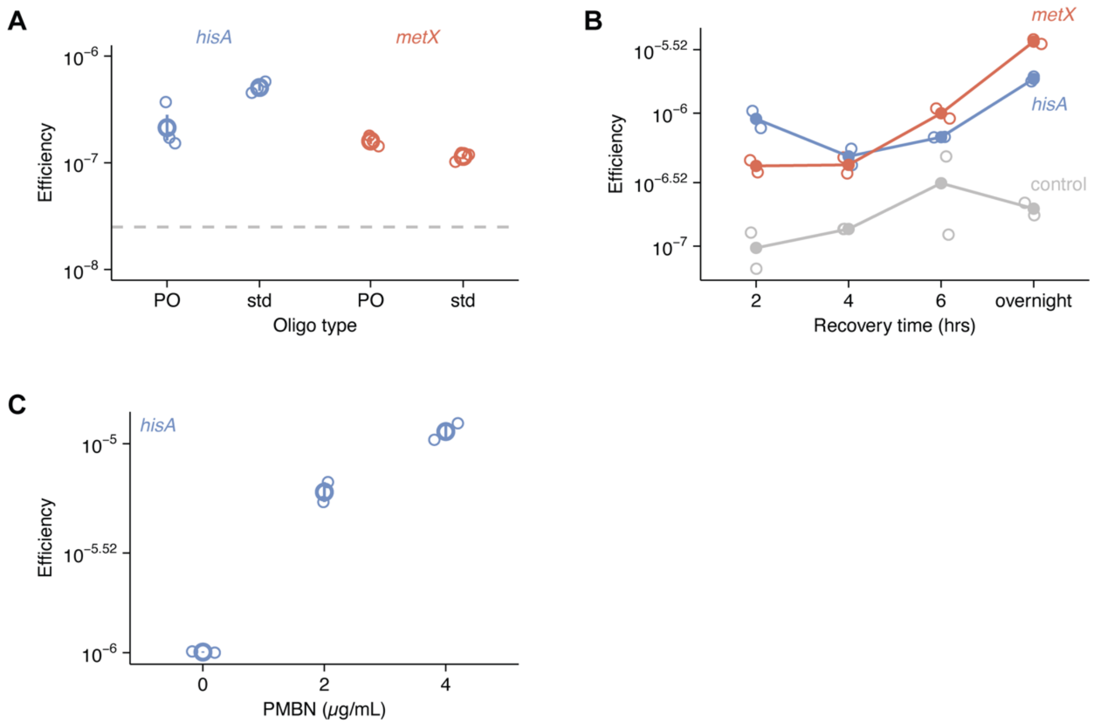

**Figure S1. ORBIT Efficiency details.** A) ORBIT efficiency for oligos with (PO) or without (std) four phosphorothioate bonds, 2x bonds at 5' and 2x bonds at 3'. Dashed line shows efficiency for the no oligo negative control. Open circles are individual transformations (n=3) and larger circles with error bars show mean and standard error. B) Efficiency for ORBIT transformations recovered for different lengths of time. Open circles are individual transformations (n=2) and filled circles show mean values. C) Efficiency after preculture with different levels of polymyxin B nonapeptide (PMBN). Point shapes are same as A (n=2).

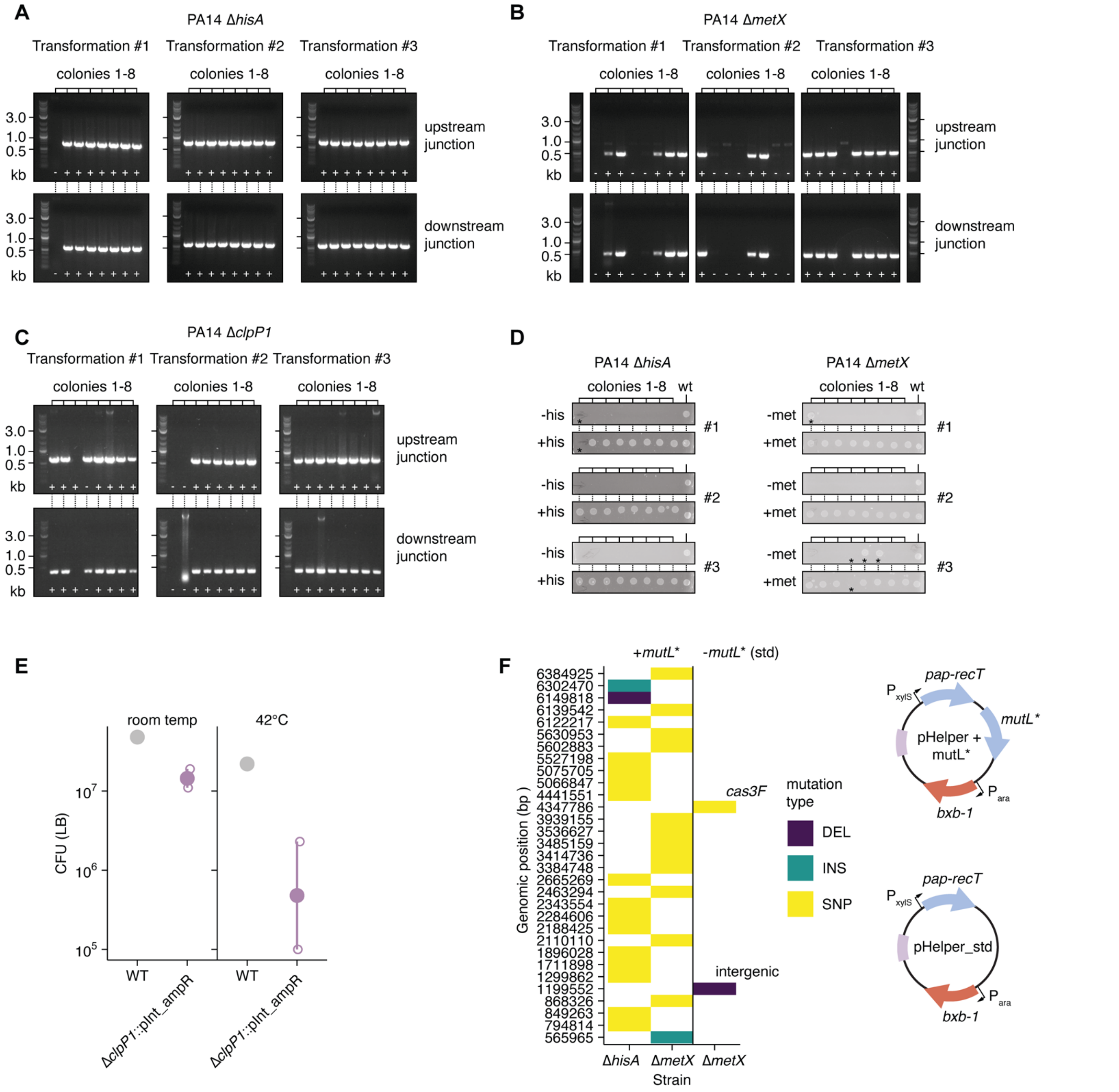

**Figure S2. Gene deletion benchmark details.** A-C) Agarose gels showing PCR genotyping assays for upstream and downstream ORBIT junctions from Δ*hisA*, Δ*metX* and Δ*clpP* PA14 strains. For three ORBIT transformations, 8 colonies each were screened. Plus symbols indicate observed bands at the correct size and accurate strain construction, while minus symbols indicate the opposite. D) Phenotype screening of colonies from A-B for growth on minimal medium with or without the expected auxotrophic amino acid. Asterisks indicate unexpected false positive or false negative results. E) Temperature sensitivity phenotypes for Δ*clpP* mutants at room temperature (~25°C) or high temperature (42°C). Open circles show replicate strains (n=1-2) and filled circles show mean values with standard error as bars. F) The genomic position of mutations from whole genome sequencing shown in Fig. 2F. Strains with a "+ *mutL**" helper plasmid are shown on the left and display many unique mutations. The only standard helper strain (std) with observed mutations is shown. The other 8 std strains showed no mutations and are not displayed in this figure.

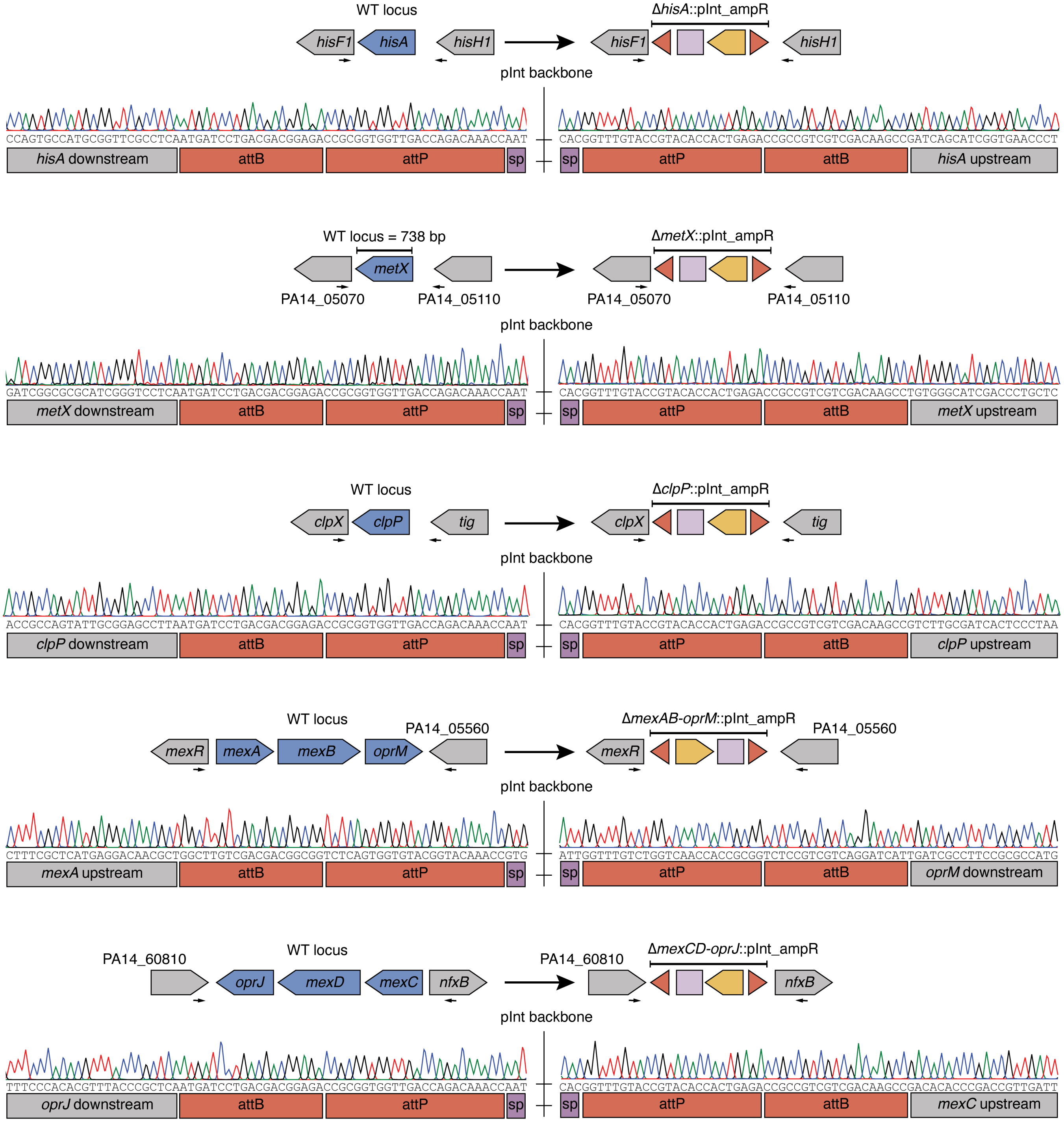

**Figure S3. Sanger sequencing of ORBIT modified loci.** The left diagram for each locus shows the original wildtype gene structure (deletion targets in blue) and the right diagram shows the structure of the ORBIT modified locus. Red annotations denote attBP sites recombined by Bxb-1. Light purple shows the RK2 origin of the integration plasmid and yellow shows the ampR marker. “sp” annotated in purple refers to the spacer sequence on the integrating plasmid. All Sanger results shown here perfectly match the reference sequences.

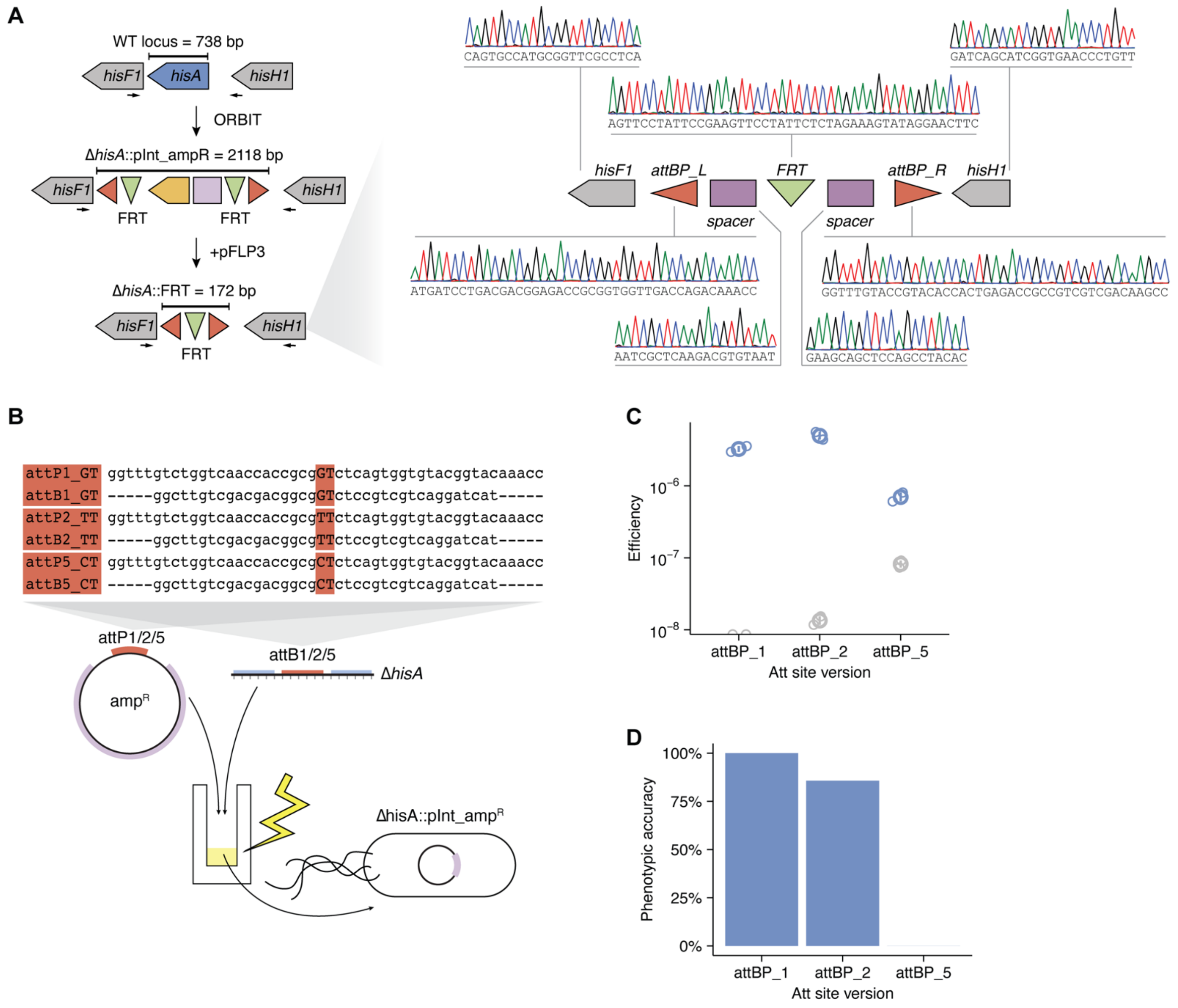

**Figure S4. Markerless mutations and alternative att sites.** A) A diagram of how ORBIT integrating plasmids are compatible with FLP-FRT excision using pFLP3. Representative perfect Sanger sequencing results of the 172 bp scar are shown for a Δ*hisA* mutant. B) Diagram of att site variants, which vary in their central dinucleotide. C-D) Efficiency and accuracy data for att site variants. Open circles represent separate transformations (n=2) and overlaid points with error bars shown mean and standard error.

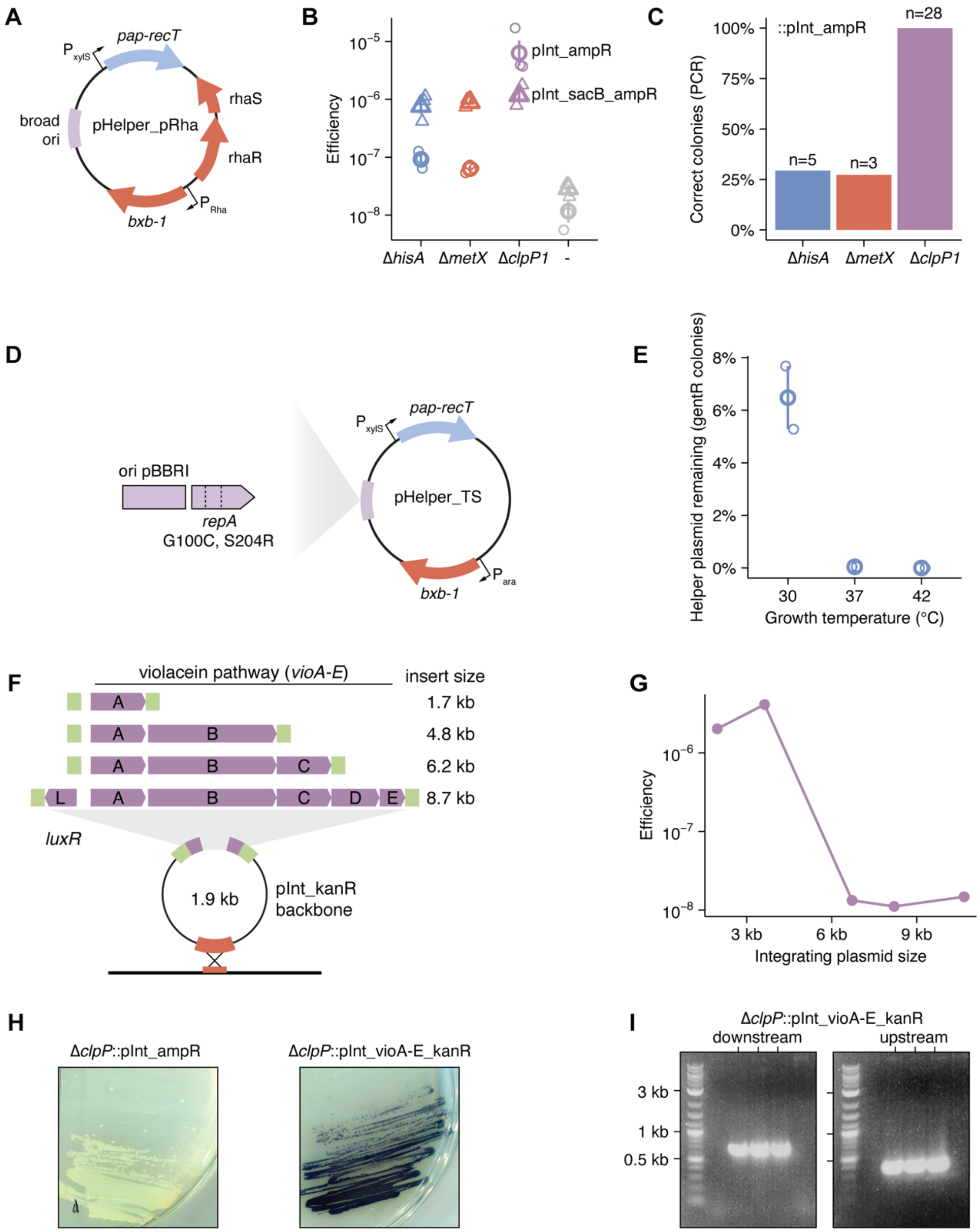

**Figure S5. Alternative helper plasmids and large integrations.** A) Diagram of pHelper_pRha with bxb-1 controlled by rhamnose induction. B-C) Efficiency and accuracy of pHelper_pRha. Open symbols show results from individual transformations and overlaid points with error bars show summary and standard error. D) Diagram of the temperature sensitive pHelper_TS with amino acid changes in the sequence of *repA*. E) The retention rate of the helper plasmid (remaining gentamicin resistance) at different plating temperatures. F-G) Diagram and efficiency results for ORBIT transformations with integrating plasmids of increasing size, ranging from the pInt_kanR backbone (1.9 kb) to a pInt with the full violacein pathway (8.7 + 1.9 = 10.6 kb). H-I) Phenotypic and molecular confirmation of the largest integration (full violacein pathway conferring purple color).

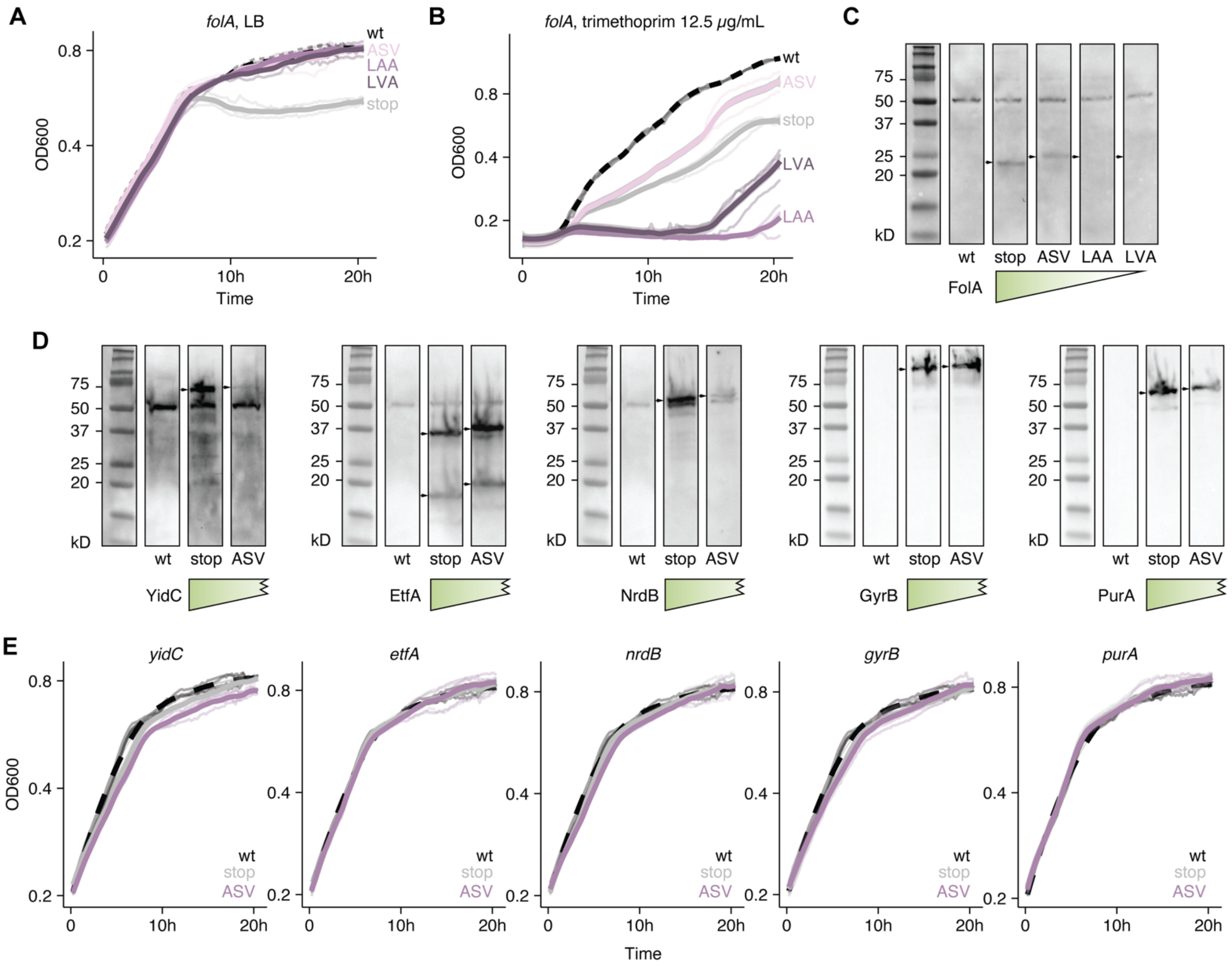

**Figure S6. Individual hypomorph phenotypes.** A) FolA degron strains growing in LB. B) FolA degron strains grown in trimethoprim (12.5 μg/mL). C) Western blots for FLAG, reporting on levels of FolA with different degron tags. D) Western blot levels of stop controls and ASV degrons for FLAG tagged gene targets (YidC, EtfA, NrdB, GyrB, and PurA). E) Growth curves (LB) for the same strains shown in D.

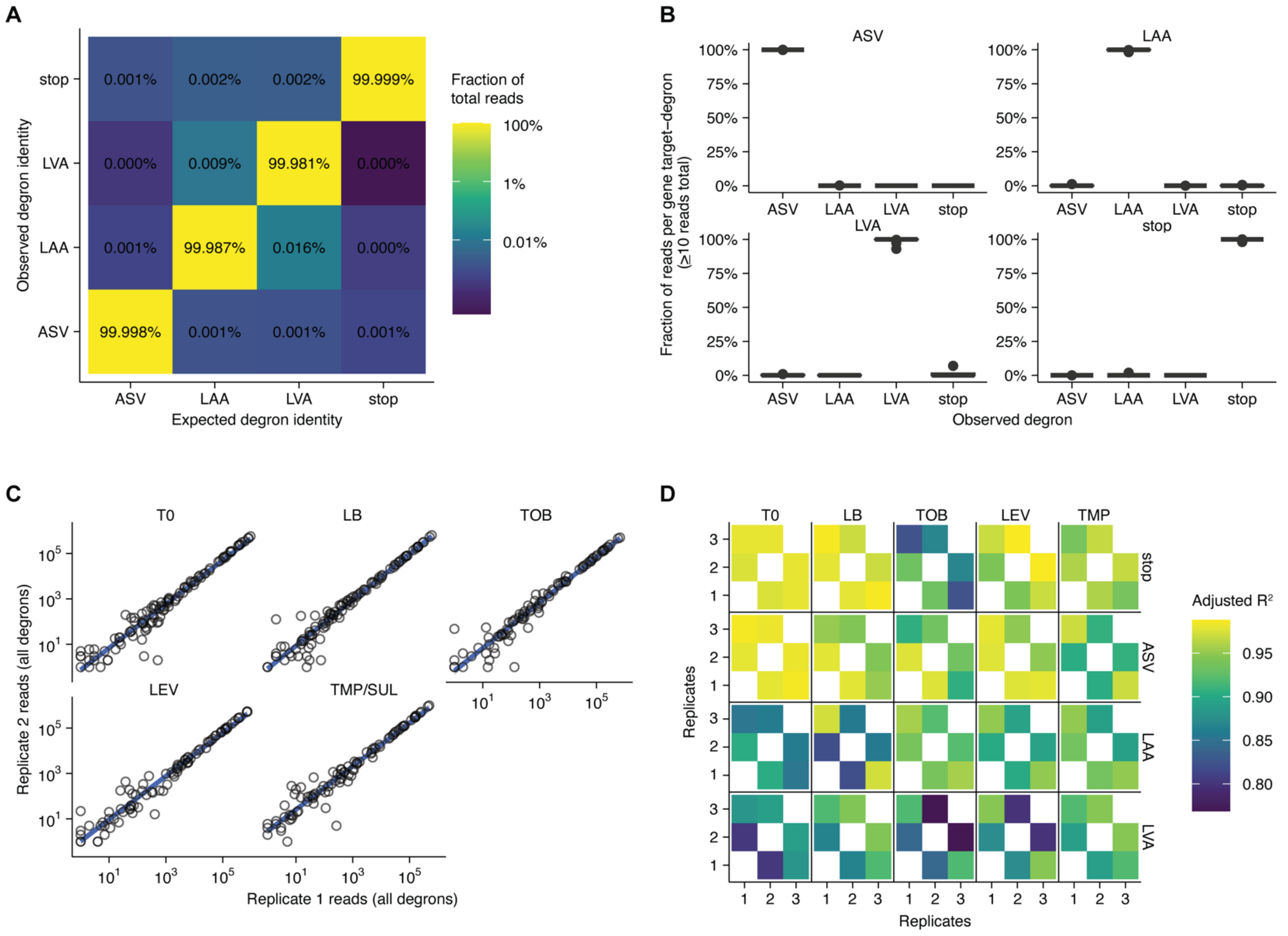

**Figure S7. Degron identifiability and reproducibility in pooled sequencing assays**. A) For all gene targets within individual degron libraries, the percentage of reads is shown that matches the expected degron identity (diagonal) or is observed as the incorrect degron sequence (off-diagonal). B) For each gene target in specific degron libraries, the distribution of degron identities is shown. C) Correlations for gene-degron strains across conditions (only replicate 1 and 2). D) $R^2$ values from linear model fits for each replicate dataset, separated by degron identity.

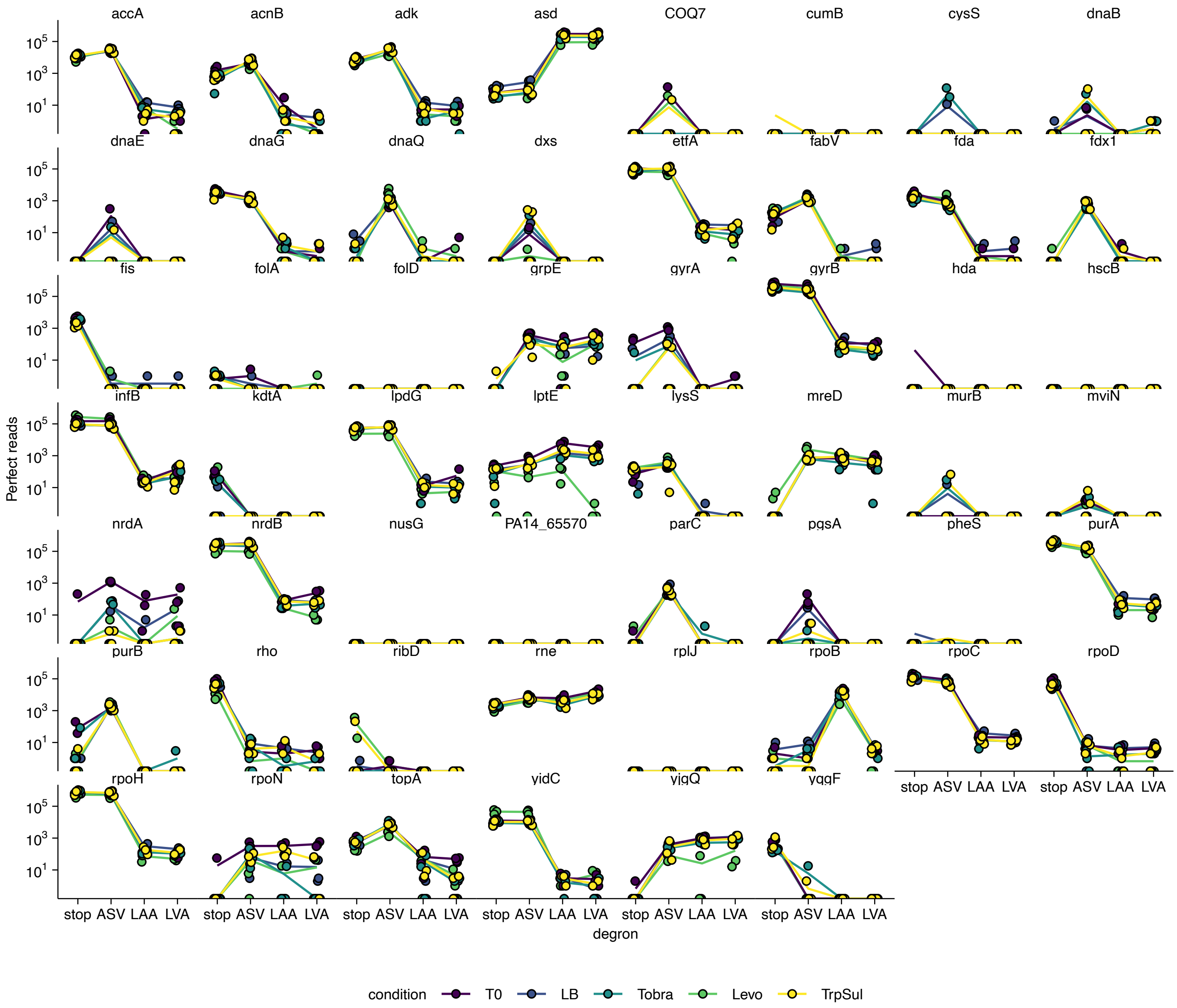

**Figure S8. Degron read count overview.** Each subpanel shows the perfect read counts for a gene target, with color indicating assay condition, and organized by increasing strength degron tags. Genes that appear to have zero reads across all conditions and degrons were detected in the initial subpool library, which is not shown here. Absent gene targets were never detected with a perfect read anywhere in the dataset.

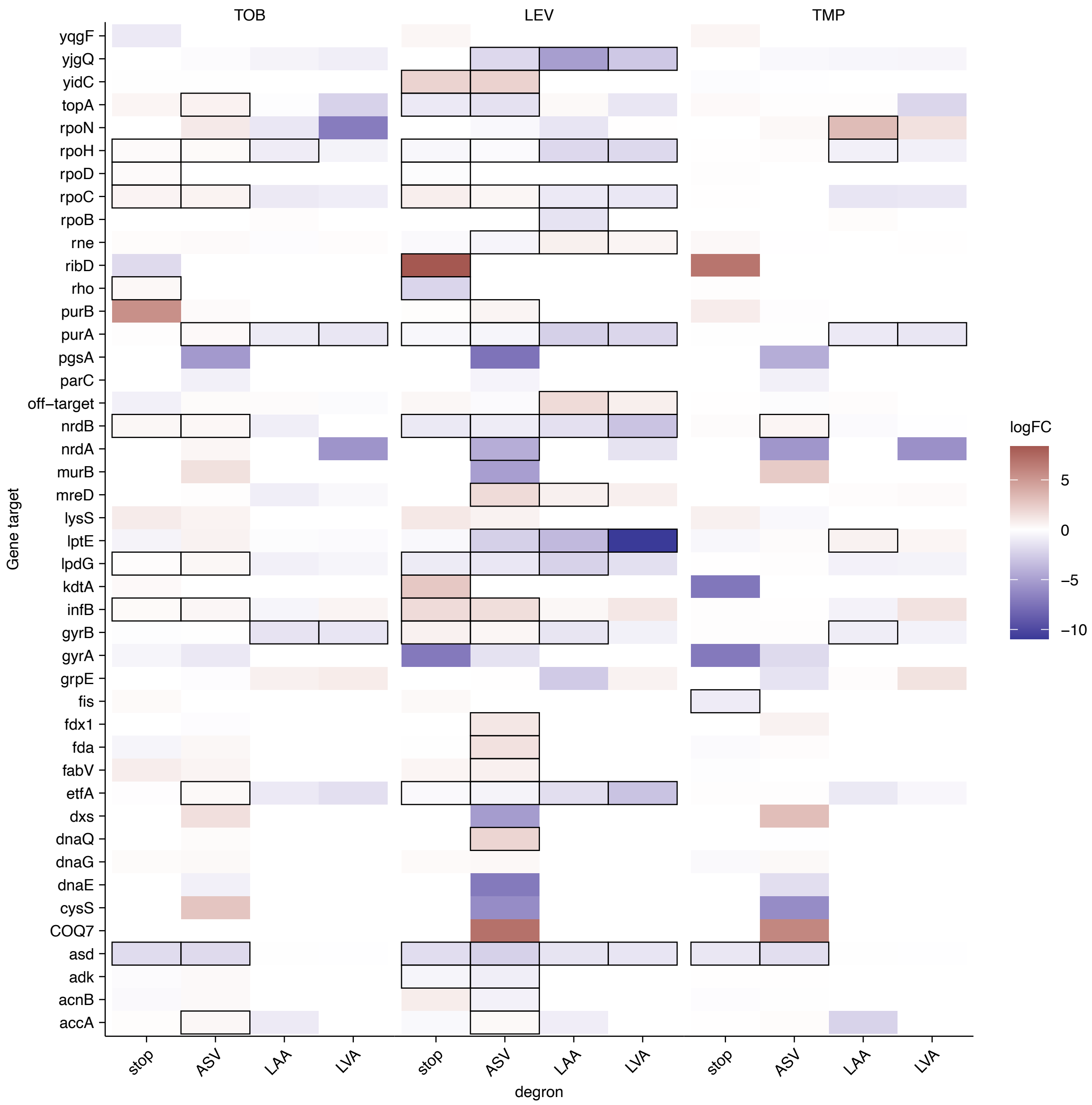

**Figure S9. Degron fitness effects across conditions**. Inferred log fold changes (logFC) are shown for each gene target and degron strain in the antibiotic conditions relative to the LB condition. Black boxes highlight results that are significant with a 10% false discovery rate. Large effects that are not significant hits are likely due to low read counts limiting statistical confidence.

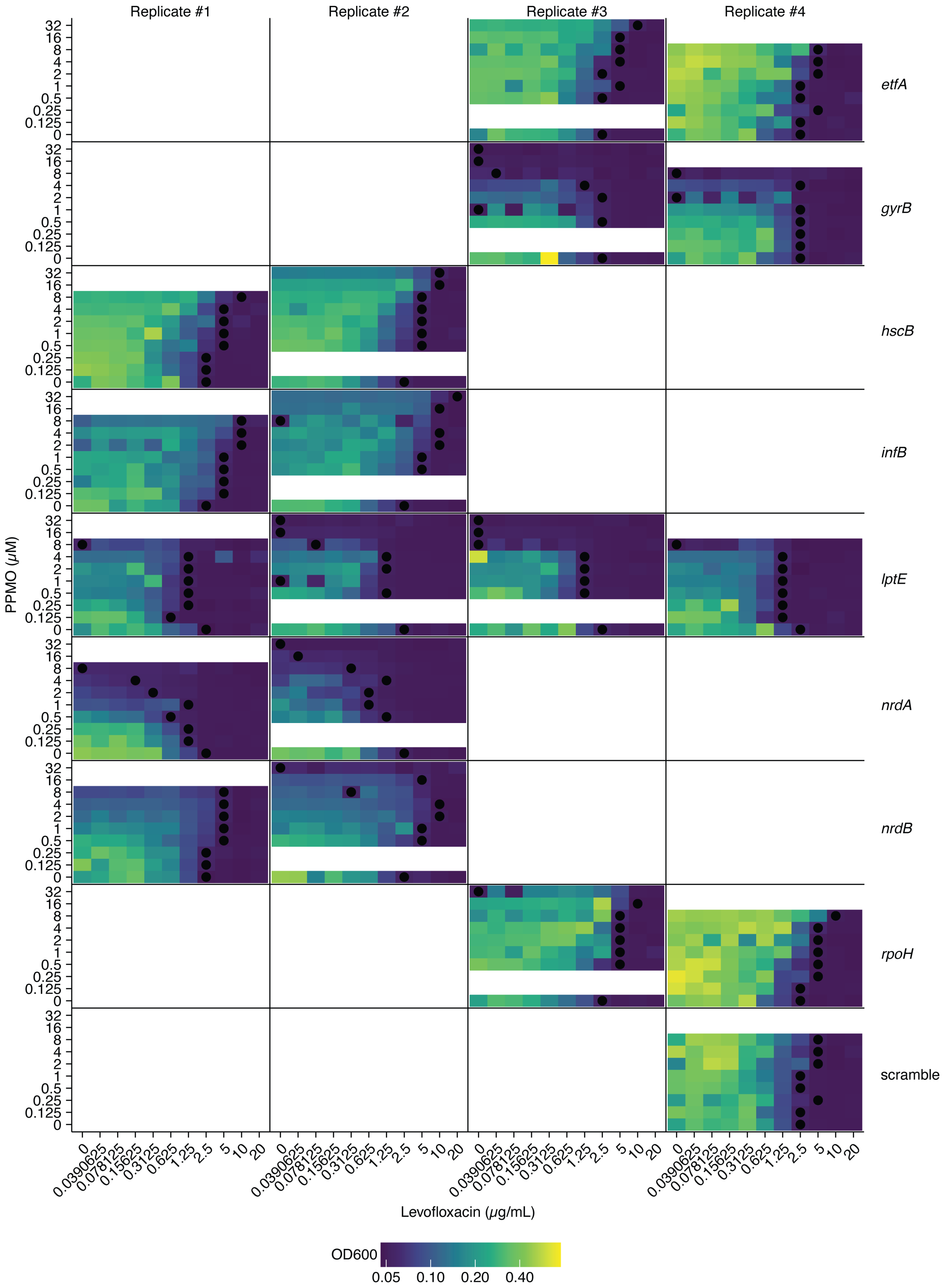

**Figure S10. PPMO checkerboard assays.** PPMOs checkerboard assay results are shown for up to four replicate experiments using different PPMOs targeting different genes shown at right. The scramble PPMO consisted of random nucleotides that did not match the Pa genome. Black dots show wells with the lowest levofloxacin concentration that did not exceed 0.06 OD units, indicating growth inhibition.

| Plasmid name | Identifier | Reference | Notes |
|---|---|---|---|
| pHelper_Pa1_std_sacB_gentR | Addgene TBD | This work | Standard helper plasmid used throughout paper. |
| pHelper_Pa1_pRha_sacB_gentR | Addgene TBD | This work | Rhamnose inducible Bxb-1 helper plasmid. |
| pHelper_Pa1_TS_sacB_gentR | Addgene TBD | This work | Temperature sensitive helper plasmid. |
| pHelper_Pa1_only_sacB_gentR | Addgene TBD | This work | RecT only helper plasmid (clean deletion). |
| pHelper_Pa1_mutL_sacB_gentR | Addgene TBD | This work | Helper plasmid with mutL* (SNPs). |
| pInt_attP1_ampR | Addgene #205296 | (15) | Basic integrating plasmid used throughout paper. |
| pInt_attP1_sacB_ampR | Addgene TBD | This work | SacB integrating plasmid used for markerless and clean modifications. |
| pInt_attP1_telR | Addgene TBD | This work | Tellurite resistant integrating plasmid. |
| pInt_attP1_ASV_ampR | Addgene TBD | This work | ASV degron integrating plasmid. |
| pInt_attP1_LAA_ampR | Addgene TBD | This work | LAA degron integrating plasmid. |
| pInt_attP1_LVA_ampR | Addgene TBD | This work | LVA degron integrating plasmid. |
| pInt_attP1_stop_ampR | Addgene TBD | This work | Stop control degron integrating plasmid. |
| pInt_attP1_3xFLAG_ASV_ampR | Addgene TBD | This work | ASV degron with FLAG integrating plasmid. |
| pInt_attP1_3xFLAG_LAA_ampR | Addgene TBD | This work | LAA degron with FLAG integrating plasmid. |
| pInt_attP1_3xFLAG_LVA_ampR | Addgene TBD | This work | LVA degron with FLAG integrating plasmid. |
| pInt_attP1_3xFLAG_stop_ampR | Addgene TBD | This work | Stop control with FLAG integrating plasmid. |
| pAra_CFP_stop_gentR | NA | This work | Replicating plasmid with CFP reporter. |
| pAra_CFP_ASV_gentR | NA | This work | Replicating plasmid with CFP and ASV degron reporter. |
| pAra_CFP_LAA_gentR | NA | This work | Replicating plasmid with CFP and ASV degron reporter. |
| pAra_CFP_AAV_gentR | NA | This work | Replicating plasmid with CFP and ASV degron reporter. |
| pAra_CFP_LVA_gentR | NA | This work | Replicating plasmid with CFP and ASV degron reporter. |
| pFLP3 | Addgene #64946 | (29) | Replicating plasmid expressing FLP for marker removal. |
| pInt_attP1_vioA_kanR | NA | (15) | Integrating plasmid with violacein pathway component. |
| pInt_attP1_vioAB_kanR | NA | (15) | Integrating plasmid with violacein pathway component. |
| pInt_attP1_vioA-C_kanR | NA | (15) | Integrating plasmid with violacein pathway component. |
| pInt_attP1_vioA-E_kanR | NA | (15) | Integrating plasmid with violacein pathway component. |

**Table S1. Plasmids used in this work.** References refer to main next numbered references.

| ID number | Genotype | Host strain | Resistance marker | Reference |
|---|---|---|---|---|
| 1 | wildtype | PA14 | none | (27) |
| 2 | wildtype | PAO1 | none | (26) |
| 3 | wildtype | TIDB #3146 | none | This work |
| 4 | wildtype | TIDB #3301 | none | This work |
| 5 | wildtype | TIDB #4041 | none | This work |
| 6 | ΔhisA::pInt_attP1_ampR | PA14 | carbenicillin | This work |
| 7 | ΔmetX::pInt_attP1_ampR | PA14 | carbenicillin | This work |
| 8 | ΔclpP::pInt_attP1_ampR | PA14 | carbenicillin | This work |
| 9 | ΔhisA::pInt_attP1_ampR | PAO1 | carbenicillin | This work |
| 10 | ΔmetX::pInt_attP1_ampR | PAO1 | carbenicillin | This work |
| 11 | ΔclpP::pInt_attP1_ampR | PAO1 | carbenicillin | This work |
| 12 | ΔmexAB-oprM::pInt_attP1_ampR | PA14 | carbenicillin | This work |
| 13 | ΔmexCD-oprJ::pInt_attP1_ampR | PA14 | carbenicillin | This work |
| 14 | ΔhisC1::pInt_attP1_ampR | PA14 | carbenicillin | This work |
| 15 | ΔphzS::pInt_attP1_ampR | PA14 | carbenicillin | This work |
| 16 | ΔhisA::attB1 | PA14 | none | This work |
| 17 | ΔmetX::attB1 | PA14 | none | This work |
| 18 | ΔhisA (clean) | PA14 | none | This work |
| 19 | ΔmetX (clean) | PA14 | none | This work |
| 20 | ΔhisA::FRT | PA14 | none | This work |
| 21 | ΔclpP::pInt_attP1_kanR | PA14 | kanamycin | This work |
| 22 | ΔclpP::pInt_vioA_attP1_kanR | PA14 | kanamycin | This work |
| 23 | ΔclpP::pInt_vioAB_attP1_kanR | PA14 | kanamycin | This work |
| 24 | ΔclpP::pInt_vioABC_attP1_kanR | PA14 | kanamycin | This work |
| 25 | ΔclpP::pInt_vioA-E_attP1_kanR | PA14 | kanamycin | This work |
| 26 | ΔclpP::pInt_attP1_telR | PA14 | tellurite | This work |
| 27 | ΔhisA::pInt_attP1_ampR | TIDB #3146 | carbenicillin | This work |
| 28 | ΔmetX::pInt_attP1_ampR | TIDB #3146 | carbenicillin | This work |
| 29 | ΔhisA::pInt_attP1_ampR | TIDB #3301 | carbenicillin | This work |
| 30 | ΔmetX::pInt_attP1_ampR | TIDB #3301 | carbenicillin | This work |
| 31 | ΔhisA::pInt_attP1_ampR | TIDB #4041 | carbenicillin | This work |
| 32 | ΔmetX::pInt_attP1_ampR | TIDB #4041 | carbenicillin | This work |
| 33 | ΔmexA::pInt_attP1_ampR | PA14 | carbenicillin | This work |
| 34 | ΔmexC::pInt_attP1_ampR | PA14 | carbenicillin | This work |
| 35 | ΔmexX::pInt_attP1_ampR | PA14 | carbenicillin | This work |
| 36 | ΔmexA::pInt_attP1_ampR | PAO1 | carbenicillin | This work |
| 37 | ΔmexC::pInt_attP1_ampR | PAO1 | carbenicillin | This work |
| 38 | ΔmexA::pInt_attP1_ampR | TIDB #3301 | carbenicillin | This work |
| 39 | ΔmexC::pInt_attP1_ampR | TIDB #3301 | carbenicillin | This work |
| 40 | ΔmexX::pInt_attP1_ampR | TIDB #3301 | carbenicillin | This work |
| 41 | ΔmexC::pInt_attP1_ampR | TIDB #3146 | carbenicillin | This work |
| 42 | ΔmexC::pInt_attP1_ampR | TIDB #4041 | carbenicillin | This work |
| 43 | hisC1::pInt_ASV_attP1_ampR | PA14 | carbenicillin | This work |
| 44 | hisC1::pInt_LAA_attP1_ampR | PA14 | carbenicillin | This work |
| 45 | hisC1::pInt_LVA_attP1_ampR | PA14 | carbenicillin | This work |
| 46 | ΔclpP (clean) | PA14 | none | This work |
| 47 | grpE::pInt_stop_attP1_ampR | PA14 | carbenicillin | This work |
| 48 | grpE::pInt_ASV_attP1_ampR | PA14 | carbenicillin | This work |
| 49 | grpE::pInt_LAA_attP1_ampR | PA14 | carbenicillin | This work |
| 50 | grpE::pInt_LVA_attP1_ampR | PA14 | carbenicillin | This work |
| 51 | grpE::pInt_3xFLAG_stop_attP1_ampR | PA14 | carbenicillin | This work |
| 52 | grpE::pInt_3xFLAG_ASV_attP1_ampR | PA14 | carbenicillin | This work |
| 53 | grpE::pInt_3xFLAG_LAA_attP1_ampR | PA14 | carbenicillin | This work |
| 54 | grpE::pInt_3xFLAG_LVA_attP1_ampR | PA14 | carbenicillin | This work |
| 55 | folA::pInt_stop_attP1_ampR | PA14 | carbenicillin | This work |
| 56 | folA::pInt_ASV_attP1_ampR | PA14 | carbenicillin | This work |
| 57 | folA::pInt_LAA_attP1_ampR | PA14 | carbenicillin | This work |

| | | | | |
|---|---|---|---|---|
| 58 | folA::pInt_LVA_attP1_ampR | PA14 | carbenicillin | This work |
| 59 | folA::pInt_3xFLAG_stop_attP1_ampR | PA14 | carbenicillin | This work |
| 60 | folA::pInt_3xFLAG_ASV_attP1_ampR | PA14 | carbenicillin | This work |
| 61 | folA::pInt_3xFLAG_LAA_attP1_ampR | PA14 | carbenicillin | This work |
| 62 | folA::pInt_3xFLAG_LVA_attP1_ampR | PA14 | carbenicillin | This work |
| 63 | dnaG::pInt_stop_attP1_ampR | PA14 | carbenicillin | This work |
| 64 | dnaG::pInt_ASV_attP1_ampR | PA14 | carbenicillin | This work |
| 65 | dnaG::pInt_3xFLAG_stop_attP1_ampR | PA14 | carbenicillin | This work |
| 66 | dnaG::pInt_3xFLAG_ASV_attP1_ampR | PA14 | carbenicillin | This work |
| 67 | yidC::pInt_3xFLAG_stop_attP1_ampR | PA14 | carbenicillin | This work |
| 68 | yidC::pInt_3xFLAG_ASV_attP1_ampR | PA14 | carbenicillin | This work |
| 69 | etfA::pInt_3xFLAG_stop_attP1_ampR | PA14 | carbenicillin | This work |
| 70 | etfA::pInt_3xFLAG_ASV_attP1_ampR | PA14 | carbenicillin | This work |
| 71 | nrdB::pInt_3xFLAG_stop_attP1_ampR | PA14 | carbenicillin | This work |
| 72 | nrdB::pInt_3xFLAG_ASV_attP1_ampR | PA14 | carbenicillin | This work |
| 73 | gyrB::pInt_3xFLAG_stop_attP1_ampR | PA14 | carbenicillin | This work |
| 74 | gyrB::pInt_3xFLAG_ASV_attP1_ampR | PA14 | carbenicillin | This work |
| 75 | purA::pInt_3xFLAG_stop_attP1_ampR | PA14 | carbenicillin | This work |
| 76 | purA::pInt_3xFLAG_ASV_attP1_ampR | PA14 | carbenicillin | This work |
| 77 | fis::pInt_3xFLAG_stop_attP1_ampR | PA14 | carbenicillin | This work |

**Table S2. Strains used in this work.** References refer to main next numbered references.
